# SARM1 detection in oligodendrocytes but not Schwann cells though *sarm1/Sarm1* deletion does not perturb CNS nor PNS myelination in zebrafish and mice

**DOI:** 10.1101/2022.12.08.519209

**Authors:** Shaline V. Fazal, Clara Mutschler, Civia Z. Chen, Mark Turmaine, Chiung-Ya Chen, Yi-Ping Hsueh, Andrea Loreto, Angeles Casillas-Bajo, Hugo Cabedo, Robin J.M. Franklin, Roger A. Barker, Kelly R. Monk, Benjamin J. Steventon, Michael P. Coleman, Jose A. Gomez-Sanchez, Peter Arthur-Farraj

## Abstract

SARM1 is a central regulator of programmed axon death and is required to initiate axon self-destruction after traumatic and toxic insults to the nervous system. Abnormal activation of this axon degeneration pathway is increasingly recognized as a contributor to human neurological disease and SARM1 knockdown or inhibition has become an attractive therapeutic strategy to preserve axon loss in a variety of disorders of the peripheral and central nervous system. Despite this, it remains unknown whether *Sarm1*/SARM1 is present in myelinating glia and whether it plays a role in myelination in the PNS or CNS. It is important to answer these questions to understand whether future therapies inhibiting SARM1 function may have unintended deleterious impacts on myelination. Here we show that *Sarm1* mRNA is present in oligodendrocytes in zebrafish but only detectable at low levels in Schwann cells in both zebrafish and mice. We find SARM1 protein is readily detectable in murine oligodendrocytes *in vitro and in vivo* and activation of endogenous SARM1 in oligodendrocytes induces cell death. In contrast, SARM1 protein is not detectable in Schwann cells and satellite glia in the adult murine nervous system. Cultured Schwann cells contain negligible functional SARM1 and are insensitive to specific SARM1 activators. Using zebrafish and mouse *Sarm1* mutants, we show that SARM1 is not required for initiation of myelination nor myelin sheath maintenance by oligodendrocytes and Schwann cells. Thus, strategies to inhibit SARM1 function in the nervous system to treat neurological disease are unlikely to perturb myelination in humans.

**Main Points:**

- SARM1 protein is detectable in oligodendrocytes but not in Schwann cells
- Oligodendrocytes but not Schwann cells die in response to endogenous SARM1 activation
- CNS nor PNS myelination, in zebrafish and mice, is hindered by loss of *sarm1/Sarm1*

## Introduction

The programmed axonal death (also termed Wallerian degeneration) pathway is becoming increasingly linked to neurological disease. Two of the most important regulators of this pathway are nicotinamide mononucleotide adenylyltransferase 2 (NMNAT2) and Sterile-α and Toll/interleukin 1receptor (TIR) motif containing protein 1 (SARM1), a member of the MyoD88 family (Gilley and Coleman, 2010; Osterloh et al., 2012; Gerdts et al., 2013; Coleman and Höke, 2020). Complete loss of function mutations in *Nmnat2* have been implicated in cases of fetal akinesia deformation sequence (FADS), where fetuses are stillborn with severe skeletal hypoplasia and hydrops fetalis, whereas partial loss of function mutations have been linked with the development of peripheral neuropathy with erythromelalgia (Huppke et al., 2019; Lukacs et al., 2019). SARM1 function is conserved in humans and the *SARM1* locus has been identified in two Genome-wide association studies (GWAS) for Amyotrophic lateral sclerosis (ALS) (Fogh et al., 2014; van Rheenen et al., 2016; Chen et al., 2021). Following this, gain of function *SARM1* mutations have been identified in sporadic ALS and other motor nerve disorders (Gilley et al., 2021; Bloom et al., 2022). Additionally, the disused rat poison, vacor, which leads to highly specific activation of SARM1, causes severe neurotoxic effects in humans. (LeWitt, 1980; Loreto et al., 2021).

Axon dysfunction and loss are hallmarks of many neurological diseases of the central (CNS) and peripheral nervous system (PNS), including Parkinson disease, traumatic brain injury, progressive multiple sclerosis, ALS and the many inherited and acquired peripheral neuropathies (Coleman and Höke, 2020). *Sarm1* deletion has shown to have significant protective effects on axon loss in animal models of traumatic brain injury, metabolic neuropathy and several models of chemotherapy induced neuropathy (Geisler et al., 2016; Henninger et al., 2016; Turkiew et al., 2017; Cheng et al., 2019; Geisler et al., 2019a; Marion et al., 2019; Bosanac et al., 2021; Gould et al., 2021b). Thus, inhibition or knockdown of SARM1 through use of pharmacological, gene therapy and antisense oligonucleotide approaches have become very attractive therapeutic strategies to trial in various neurological diseases (Geisler et al., 2019b; Coleman and Höke, 2020; Krauss et al., 2020; Arthur-Farraj and Coleman, 2021; Bosanac et al., 2021; Gould et al., 2021a; Merlini et al., 2022).

SARM1 is abundant in the nervous system but is not found in most other tissues, including heart, kidney, liver, lung, skeletal muscle, spleen or thymus. (Kim et al., 2007; Chen et al., 2011). While neurons have been shown to have high levels of SARM1 it is unknown whether it is also present in myelinating glia in the PNS or CNS. Furthermore, it remains undetermined, due to the lack of in-depth quantitative studies on myelination, whether SARM1 has any role in regulating oligodendrocyte or Schwann cell myelination or myelin maintenance in either a cell autonomous or non-cell autonomous fashion. This is important to know as it could have adverse implications for the use of therapies aimed to treat various neurological disorders through inhibition of SARM1 function.

In this study, we use mice and larval zebrafish to demonstrate that *Sarm*1/*sarm1* mRNA is present at low levels in developing Schwann cells and oligodendrocytes. In the adult murine nervous system SARM1 protein is detectable in oligodendrocytes but not in Schwann cells and satellite glia. In cultured cells, use of the specific SARM1 activators, vacor and 3-acetylpyridine (3-AP) confirms that Schwann cells contain negligible amounts of SARM1 whereas cultured oligodendrocytes contain functionally relevant levels of SARM1 protein. Furthermore, we show that, in the absence of SARM1/Sarm1, myelination in the PNS and CNS is initiated in a timely fashion in both mice and larval zebrafish and that PNS and CNS myelin maintenance in adult mice is unaffected.

## Results

### SARM1 protein is present in oligodendrocytes but not in Schwann cells nor satellite glia

In order to investigate the presence of SARM1 protein in various cell types in the murine nervous system we used a validated polyclonal antibody generated by C.-Y. Chen et al., 2011 in transverse cryosections of adult (P60) tibial and optic nerve. As expected, we found that SARM1 expression colocalized with the axonal markers neurofilament and beta-III tubulin in sciatic and optic nerve respectively (Fig.1A and B). Additionally, we found that SARM1 protein was present ubiquitously, at high levels, in large and small dorsal root ganglion (DRG) neuron cell bodies, *in vivo*, compared to lower levels within their axons (Fig. 1C). Importantly, we saw no staining in control sections without primary antibody nor in *Sarm1* knockout (KO) DRG neurons (Sup Fig. 1A and B). To test if there were detectable levels of SARM1 protein in myelinating and non-myelinating Schwann cells, satellite glia and oligodendrocytes we performed SARM1 and SOX10 immunohistochemistry in cryosections from tibial nerves, DRGs and optic nerves. In PNS tissue we found no colocalization in multiple sections from 3 separate non-littermate animals (Fig. 1D-E). However, in optic nerve sections SARM1 protein appeared present in a perinuclear staining pattern around SOX10 positive nuclei (Fig. 1F). To confirm this, we performed SARM1 immunohistochemistry in optic nerves of two mutant mouse lines where oligodendrocytes are labelled with tdTomato; either *Plp1-cre/ERT2* or *Sox10-cre* mouse lines bred with homozygous *Rosa26 stopflox tdtomato* mice. In both lines, tdTomato positive cells contained SARM1 immunolabelling, including in single confocal slices (Fig. 1G and H). Thus, SARM1 protein appears restricted to neuronal populations in the PNS and is not detectable in adult murine Schwann cells and satellite glia. However, in the adult mammalian CNS, SARM1 protein is readily detectable in oligodendrocytes.

**Figure 1.**
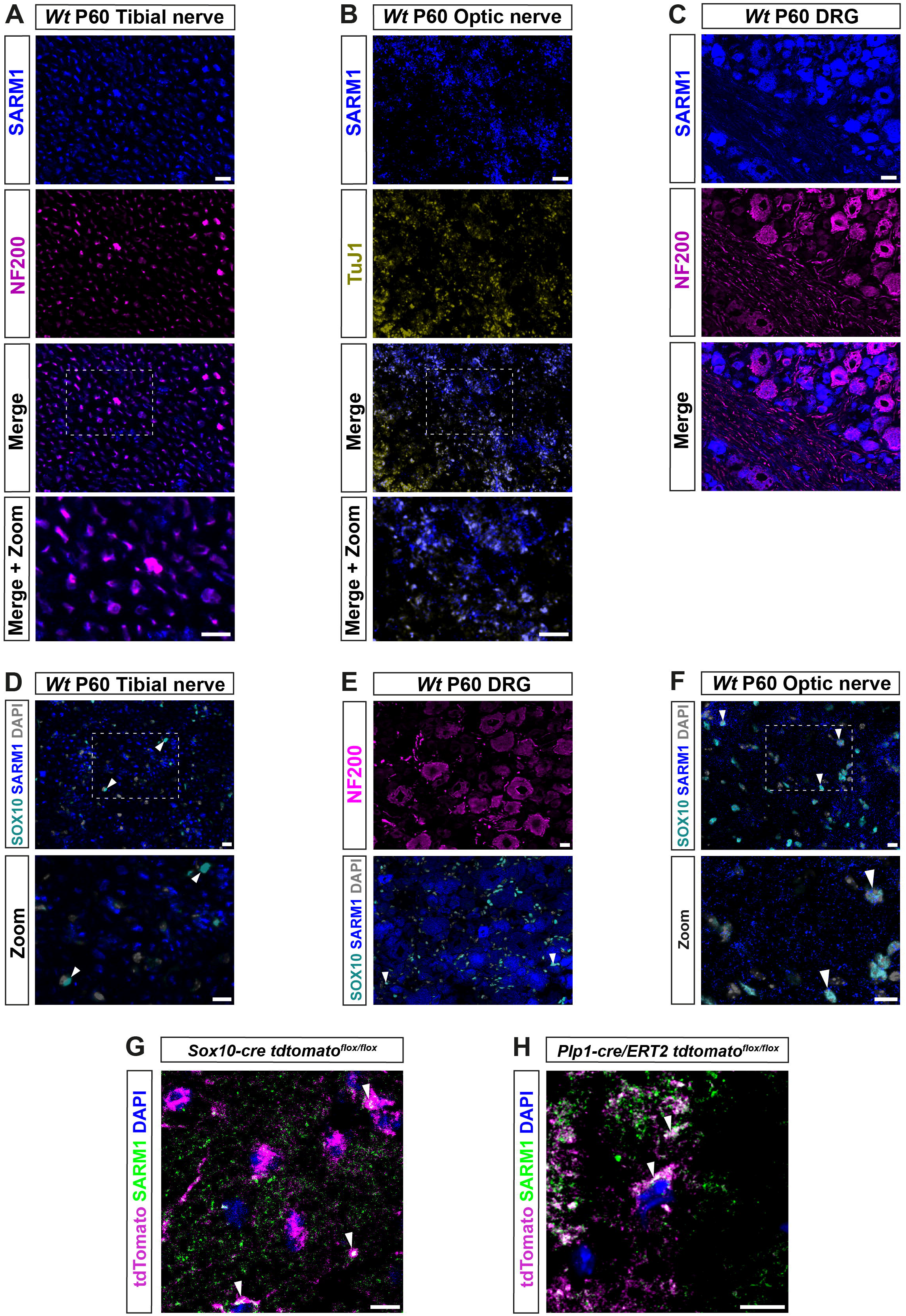
SARM1 protein is detectable in CNS myelinating glia but not PNS glia. A) Representative immunofluorescence images of transverse sections of *Wt* mouse tibial nerves to show co-localization of SARM1 (blue) and NF-200 (neurofilament light chain; magenta) in axons. Zoomed area is represented by the white bounding box. Scale bars 10 μm. B) Representative immunofluorescence images of transverse sections of *Wt* mouse optic nerves to show expression of SARM1 (blue) in TUJ1 (Anti-Beta III Tubulin; yellow) positive axons. Zoomed area is represented by the white bounding box. Scale bars 10 μm. C) Representative immunofluorescence images of transverse sections of *Wt* mouse DRGs to show co-localization of SARM1 (blue) and NF-200 (magenta) positive neurons. Scale bar 25 μm. D) Representative immunofluorescence images of transverse sections of *Wt* and *Sarm1* KO mouse tibial nerves showing SARM1 (blue) and SOX10 (cyan) expression. White arrowheads show co-localization of DAPI (gray) and SOX10 (cyan), but no expression of SARM1 (blue). Zoomed areas are represented by the white bounding box. Scale bar 10 μm. E) Representative immunofluorescence images of transverse sections of *Wt* and *Sarm1* KO mouse DRGs showing SARM1 (blue) and SOX10 (cyan) expression. White arrowheads point to SOX10 (cyan) positive satellite glia and DAPI (gray), with no colocalization of SARM1 (blue). Scale bar 10 μm. F) Representative immunofluorescence images of transverse sections of *Wt* and *Sarm1* KO mouse optic nerves showing SARM1 (blue) and SOX10 (cyan) expression. White arrowheads point to SOX10 (cyan) positive oligodendrocytes and DAPI (gray), with perinuclear SARM1 staining (blue). Scale bar 10 μm. Zoomed areas are represented by the white bounding box. Scale bar 10 μm. All experiments n=3 (3 animals, 2 nerves per animal, 5 sections per nerve). G) Maximum projection of tdTomato positive oligodendrocytes in P60 optic nerves of *Sox10-cre Rosa26 stopflox tdtomato* (*Sox10-cre tdtomato*^*flox/flox*^) positive for SARM1 immunolabelling (white arrowhead). Scale bar 10 μm. H) Single confocal z plane of tdTomato positive oligodendrocytes in P60 optic nerves of *Plp1-cre/ERT2 Rosa26 stopflox tdtomato* (*Plp1-cre/ERT2 tdtomato*^*flox/flox*^) positive for SARM1 immunolabelling (white arrowhead). Scale bar 10 μm.

To identify if *sarm1* mRNA is expressed in myelinating glia in zebrafish, we used third generation in situ hybridization chain reaction (HCR), in five-day post fertilization (5dpf) zebrafish larvae, where both oligodendrocytes and Schwann cells, in the CNS and PNS, respectively, are initiating myelination (D’Rozario et al., 2017; Choi et al., 2018). We used HCR probes targeted to *sarm1* and *sox10* to label cells of the oligodendrocyte and Schwann cell lineage. In the posterior lateral line nerve (PLLn) we saw strong expression of *sox10* but little *sarm1* expression, though importantly we saw no *sarm1* signal when antisense probes were omitted (Fig. 2A; Sup Fig. 2A and B). To investigate whether *sarm1* and *sox10* were co-expressed in the same cells in the PNS, we examined individual confocal slices (616nm optical thickness) and found that we could only see co-expression of both markers in nuclei along the PLLn infrequently (Fig. 2B). Specifically, we found that 22.4%+/-3.161 (n=5 larvae, 5 sections per larvae) of Schwann cells, in the PLLn, express a low level of *sarm1*. In the spinal cord, while we saw *sox10* expression mainly concentrated in the ventral and dorsal spinal cord tracts, we visualized *sarm1* expression more diffusely throughout the whole spinal cord (Fig. 2C). In single confocal slices, the majority of *sox10* positive cells also expressed *sarm1* (Fig. 2D).

**Figure 2.**
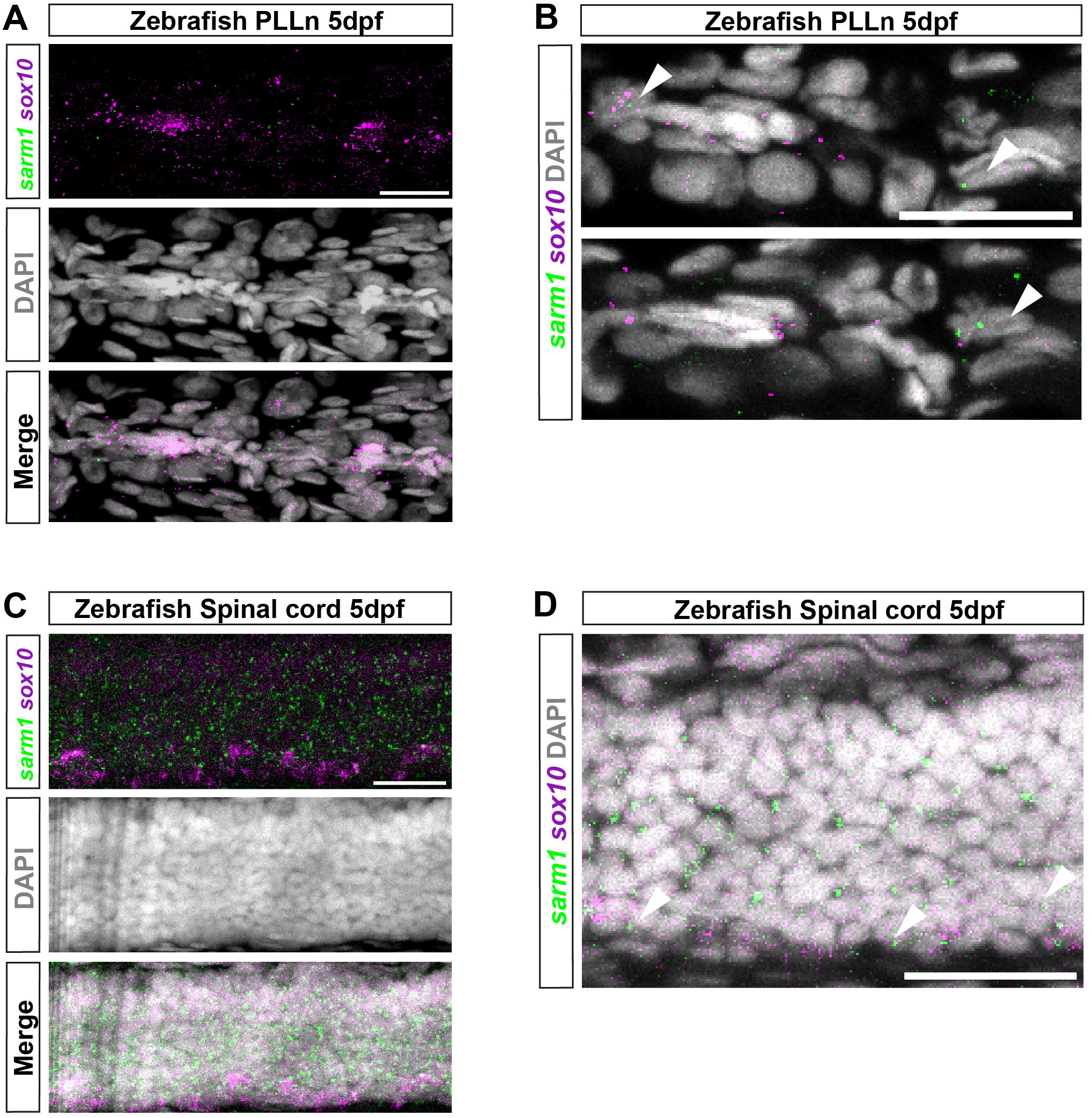
*sarm1* expression in zebrafish larvae PLLn and spinal cord. A) MAX projection, lateral view of PLLn of a 5dpf larva demonstrating HCR *in situ* labelling for *sox10* and *sarm1* mRNA and DAPI staining to mark nuclei. B) Single confocal z plane, 616 nm optical thickness, lateral view of PLLn of a 5dpf larva. Arrowheads mark nuclei, labelled with DAPI, positive for *sox10* and *sarm1* mRNA. C) MAX projection, lateral view of spinal cord of a 5dpf larva demonstrating HCR *in situ* labelling for *sox10* and *sarm1* mRNA and DAPI staining to mark nuclei. D) Single confocal z plane, 616 nm optical thickness, lateral view of spinal cord of a 5dpf larva. Arrowheads mark nuclei positive for *sox10* and *sarm1* mRNA. For all images: *sox10* mRNA (magenta) and *sarm1* mRNA (green), nuclei labelled with DAPI (gray), scale bar 25 μm. All experiments n=7 biological replicates.

In combination, these results are consistent with the view that peripheral glia such as Schwann cells and satellite glia do not contain detectable levels of SARM1, whereas oligodendrocytes in the CNS express substantial levels of *sarm1*/SARM1 in fish and mice.

### Cultured oligodendrocytes, but not Schwann cells, have detectable SARM1 protein and are vulnerable to specific, SARM1 activator-induced cell death

Since we were unable to identify SARM1 protein in Schwann cells *in vivo* but we did find that a proportion of zebrafish Schwann cells express low levels of *Sarm1* mRNA, we wanted to further explore whether Schwann cells may contain small amounts of functional SARM1 protein below the threshold of antibody detection. Firstly, we confirmed that cultured mouse Schwann cells also express very low levels of *Sarm1* mRNA, similar to our finding in zebrafish larvae (Sup. Fig. 3A). Additionally, we were unable to detect SARM1 protein in Schwann cell cultures (Sup. Fig. 3B)

To test if Schwann cells have any functional SARM1 protein, we treated cultures of freshly isolated mouse Schwann cells with two separate SARM1 activators, vacor and 3-AP. Addition of high dose vacor (100 μM) or 3-AP (250 μM) to neuronal cultures induces rapid SARM1 activation, profound Nicotinamide adenine dinucleotide (NAD^+^) depletion and axonal degeneration and cell death within hours (Loreto et al., 2021; Wu et al., 2021). We found that mouse Schwann cells were completely insensitive to induction of cell death by both vacor and 3-AP treatment at high doses for long periods of time (up to 72 hours) (Fig. 3A and Sup Fig. 4A). Furthermore, Schwann cells treated with vacor or 3-AP for 72 hours showed no reduction in intracellular NAD^+^ levels, unlike in Human embryonic kidney cells, which express low levels of SARM1 protein (Fig. 3B and Sup Fig. 4B). In contrast, we found that treatment of rat oligodendrocyte cultures with vacor induced cell death within hours, and complete loss of cultures after 72 hours of treatment (Fig. 3C and D). Predictably, when we immunolabelled oligodendrocyte cultures we found the presence of detectable levels of SARM1 using two different antibodies, in line with our *in vivo* findings (Fig. 3E and Sup Fig. 4C). Thus, cultured Schwann cells, in contrast to oligodendrocytes, do not contain sufficient endogenous SARM1 protein to induce either cell death or a decline in cellular NAD^+^ levels.

**Figure 3.**
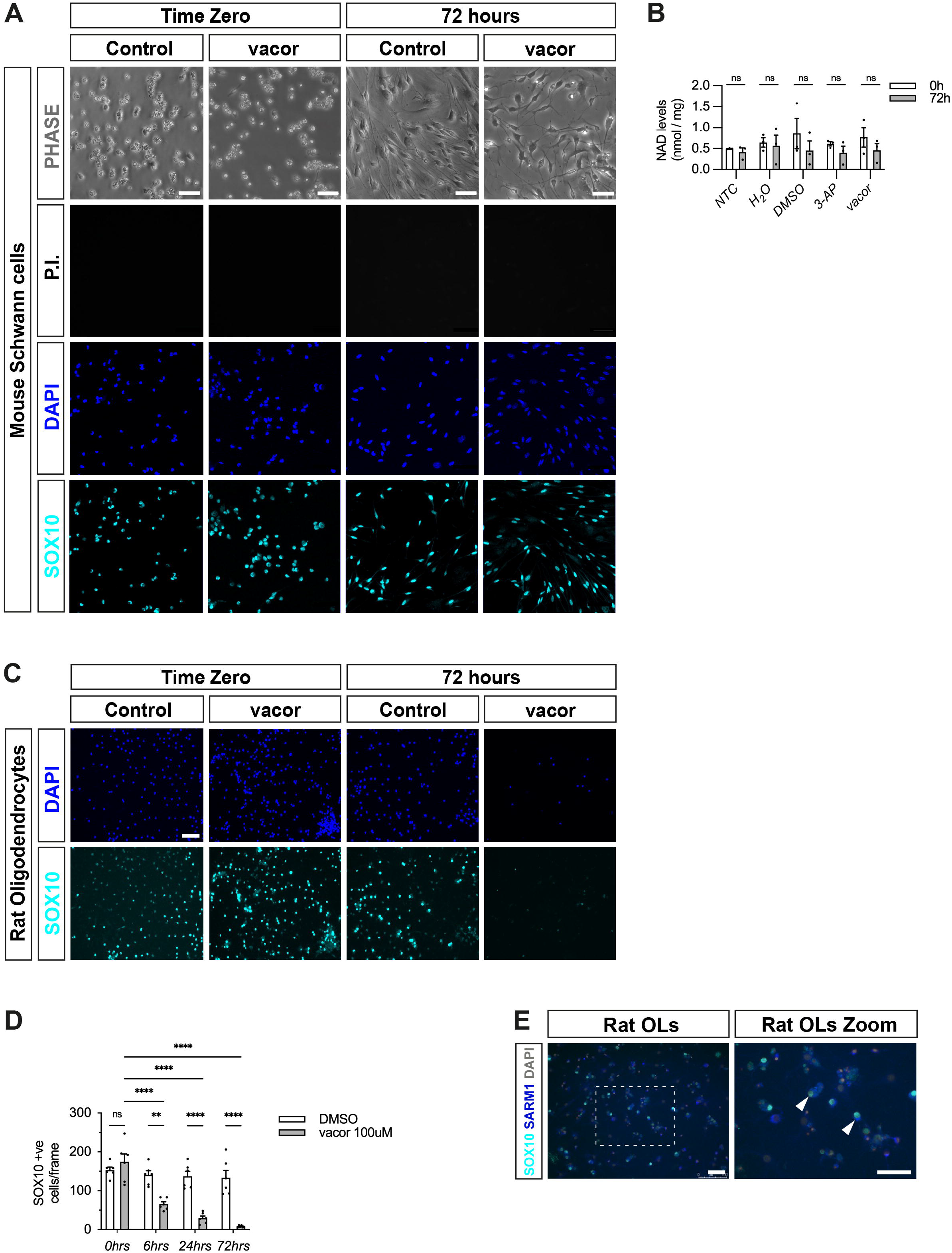
Cultured oligodendrocytes, but not Schwann cells, are sensitive to specific SARM1 activators. A) Dissociated and freshly plated P2 mouse Schwann cells cultured in DMSO or 100 μM vacor for 72 hours show no cell death, judged by propidium iodide (P.I) and DAPI nuclear staining and SOX10 immunocytochemistry (n=3: two-four mice, each from three litters producing three separate cultures). Scale bar 25 μm. B) Intracellular NAD^+^ levels do not change in Schwann cells when treated for 72 hours with either 250 μM 3-AP or 100 μM vacor compared to control conditions. C) Rat oligodendrocytes cultured in DMSO or 100 μM vacor for 72 hours demonstrate substantial cell death judged by DAPI nuclear staining and SOX10 immunocytochemistry (n=3: one rat from three litters producing three separate cultures). Scale bar 50 μm. D) Quantification of SOX10 positive oligodendrocyte cell survival in response to 100 μM vacor treatment compared to DMSO control cultures. E) Cultured rat oligodendrocytes (OLs; SOX10 positive) express SARM1 protein using a polyclonal anti-SARM1 antibody generated by Yi-Ping Hsueh. Zoomed in area (indicated by dashed box). Arrowheads indicate SOX10 positive SARM1 positive cells. Scale bars 50 μm.

### Initiation of CNS and PNS myelination proceeds normally in *Sarm1* mutant zebrafish larvae

Since inhibition of SARM1 function is being proposed to treat neurological disease, it is crucial to identify whether loss of SARM1 leads to any disruption of vertebrate CNS or PNS myelination in a cell autonomous or non-cell autonomous fashion. This is especially important given we have now shown that oligodendrocytes contain functional SARM1 protein *in vitro* and *in vivo*. To date, no studies have performed an in-depth quantitative analysis of CNS and PNS myelination in the absence of SARM1 function. To test whether Sarm1 is required for initiation of myelination in zebrafish we assessed myelination and myelin gene expression in the spinal cord and PLLn in *wild-type (wt)* and *sarm1*^*SA11193/SA11193*^ (mutant) larvae at 5dpf. The *sarm1*^*SA11193*^ mutants fish harbour a point mutation (C>A) in exon 2, which introduces a premature stop codon (Kettleborough et al., 2013). *sarm1*^*SA11193*^ mutants are viable, fertile and appear morphologically indistinct from *wt* fish (data not shown). To confirm that *sarm1* mutants behave functionally like *sarm1* null animals, which have substantially delayed axonal degeneration after traumatic injury, we injected *wt* and homozygous mutant fish with a *neuroD:tdTomato* DNA construct at the one-celled zygote stage. We then performed 2-photon laser axotomy of tdTomato labelled PLLn neurons at 4dpf and live imaged the axons distal to the injury site (Sup Fig. 5A). We found that while *wt* axons started to degenerate between 2hr 40 minutes and 3 hour 20 minutes (n=7), axons in *sarm1* mutant fish remained intact even 26 hours after axotomy, which was the limit of our imaging abilities due to UK home office regulations (Sup Fig. 5B). This confirmed that *sarm1*^*SA11193*^ mutant zebrafish phenotypically behave like *Sarm1* null mice and are similar to a Crispr-Cas9 generated *sarm1* null mutant zebrafish (Tian et al., 2020).

To assess PNS and CNS myelination, we next crossed *sarm1*^*SA11193*^ mutants with *Tg(mbp:EGFP-CAAX)* zebrafish that express membrane bound GFP under the myelin basic protein (*mbp*) promoter (Almeida et al., 2011). We observed normal formation of long GFP-labelled myelin segments in the dorsal and ventral spinal cord as well as PLLn in both 5dpf *sarm1*^*SA11193*^ mutants and *wt* fish and quantification showed no differences in PLLn and spinal cord myelination (Fig. 4A-F).

**Figure 4.**
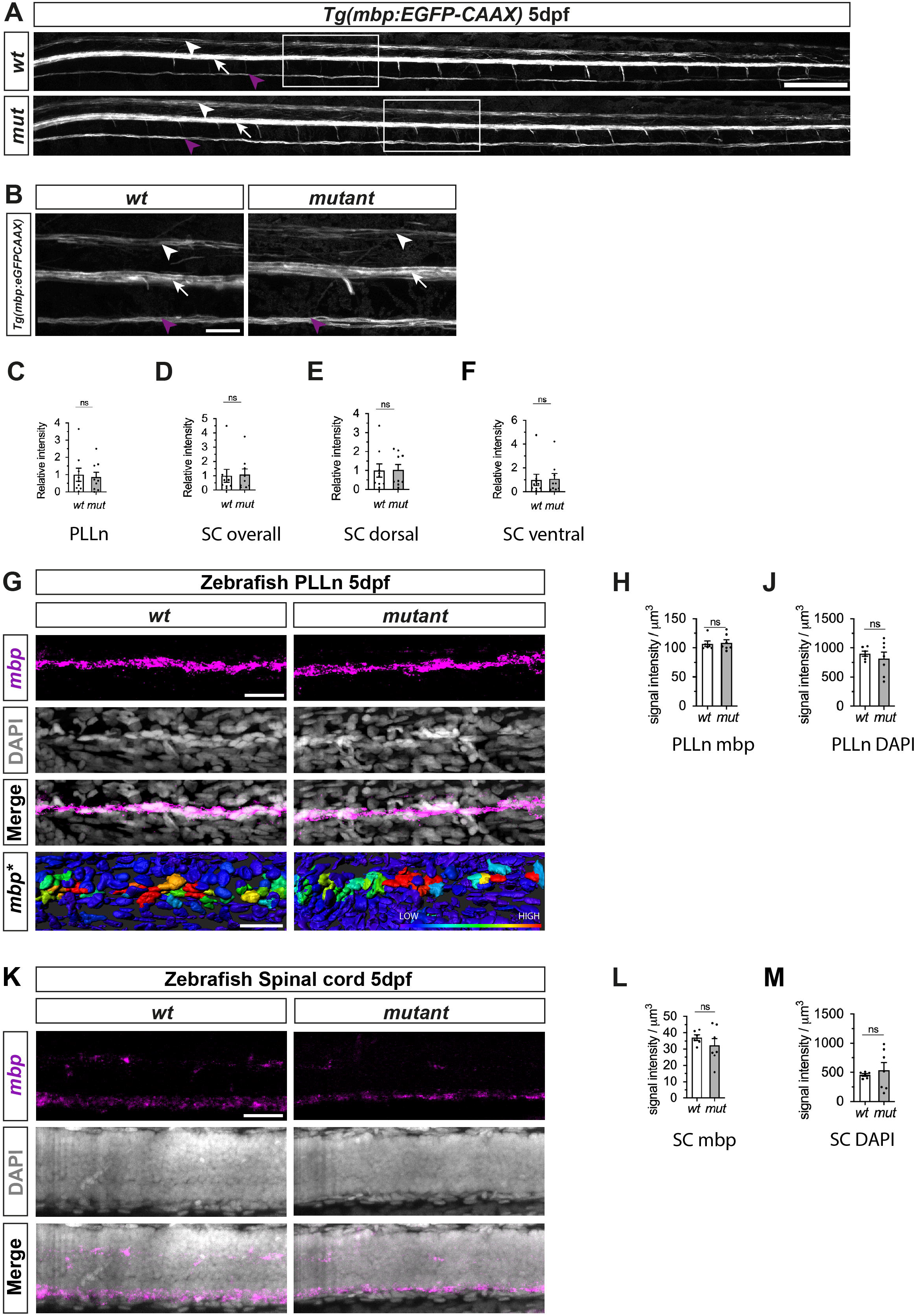
Myelination of PLLn and spinal cord is unaffected by absence of functional Sarm1 in zebrafish. A) Max projection of the lateral view of *wild-type* (*wt*) *Tg[mbp:EGFP-CAAX]* and *sarm1*^*SA11193/SA11193*^ (*sarm1 mutant, mut*) *Tg[mbp:EGFP-CAAX]* 5dpf larva. Arrowhead (white) delineates the dorsal spinal cord; Arrow (white) marks the ventral spinal cord; Arrow (magenta) pinpoints the PLLn. Scale bar 100 μm. B) Zoomed in lateral view of spinal cord and PLLn (area indicated by boxes in A). Arrowhead (white) delineates the dorsal spinal cord; Arrow (white) marks the ventral spinal cord; Arrow (magenta). Scale bar 25 μm. C) Relative intensity of GFP in PLLn of *Tg[mbp:EGFP-CAAX] wt* and *sarm1 mutant* (*mut*) (n=9; p=08633). D) Relative intensity of GFP in the dorsal and ventral spinal cord combined of *Tg[mbp:EGFP-CAAX] wt* and *sarm1 mutant* (*mut*) (n=9; p=08633). E) Relative intensity of GFP in the dorsal spinal cord of *Tg[mbp: EGFP-CAAX] wt* and *sarm1 mutant* (*mut*) (n=9; p=09314). F) Relative intensity of GFP in the ventral spinal cord of *Tg[mbp:EGFP-CAAX] wt* and *sarm1 mutant* (*mut*) (n=9; p=08633). G) MAX projection, lateral view of PLLn of *wt* and *sarm1 mutant* (*mutant*) 5dpf larvae showing *mbp* (magenta) mRNA expression. Nuclei labelled with DAPI (gray). Scale bar 25 μm. Heatmap of intensity of nuclear *mbp* mRNA expression (*mbp\**) in PLLn of *wt* and *sarm1 mutant* (*mut*) 5dpf larvae. Scale bar 20 μm H) Quantification of *mbp* mRNA signal intensity/um in PLLn of *wt* and *sarm1 mutant* (*mut*) 5dpf larvae (WT n=6; MUT n=7; p=0.8357). J) Quantification of DAPI signal intensity/um in PLLn of *wt* and *sarm1 mutant* (*mut*) 5dpf larvae (WT n=6; MUT n=7; p=0.7308). K) MAX projection, lateral view of spinal cord of *wt* and *sarm1 mutant* (*mut*) 5dpf larvae showing *mbp* (magenta) mRNA expression. Nuclei labelled with DAPI (gray). Scale bar 25 μm. L) Quantification of *mbp* mRNA signal intensity/μm in dorsal and ventral spinal cord combined of *wt* and *sarm1 mutant* (*mut*) 5dpf larvae (*wt* n=6; *mut* n=7; p=0.4452). M) Quantification of DAPI signal intensity/um in spinal cord of *wt* and *sarm1 mutant* (*mut*) 5dpf larvae (*wt* n=6; *mut* n=7; p=0.7308).

In order to quantify myelin gene expression between *wt* and *sarm1* mutant zebrafish, we performed *mbp* HCR. We found that there were equivalent levels of *mbp* expression in the spinal cord and in the PLLn in *wt* and *sarm1*^*SA11193*^ mutant fish and importantly, there were no differences in cell numbers in either structure between genotypes (Fig. 4G-M).

Collectively these experiments demonstrate that myelination proceeds normally in both the PLLn and spinal cord in zebrafish larvae in the absence of Sarm1 function.

### PNS myelination and myelin maintenance are normal in the *Sarm1* KO mouse

Given that zebrafish embryos appear to myelinate normally without functional Sarm1, we next tested whether PNS myelination in *Sarm1* KO mice is initiated on time. PNS myelination starts shortly after birth in murine peripheral nerves (Jessen and Mirsky, 2005). We looked at postnatal (P) day 2 transverse sections through the tibial nerve by transmission electron microscopy (EM) and noted that while there was a non-significant trend towards slightly more axons and Schwann cells in the *Sarm1* KO nerves compared to *Wt* (n=5), there was no difference in the proportion of large caliber axons that were myelinated (Fig. 5A-H).

**Figure 5.**
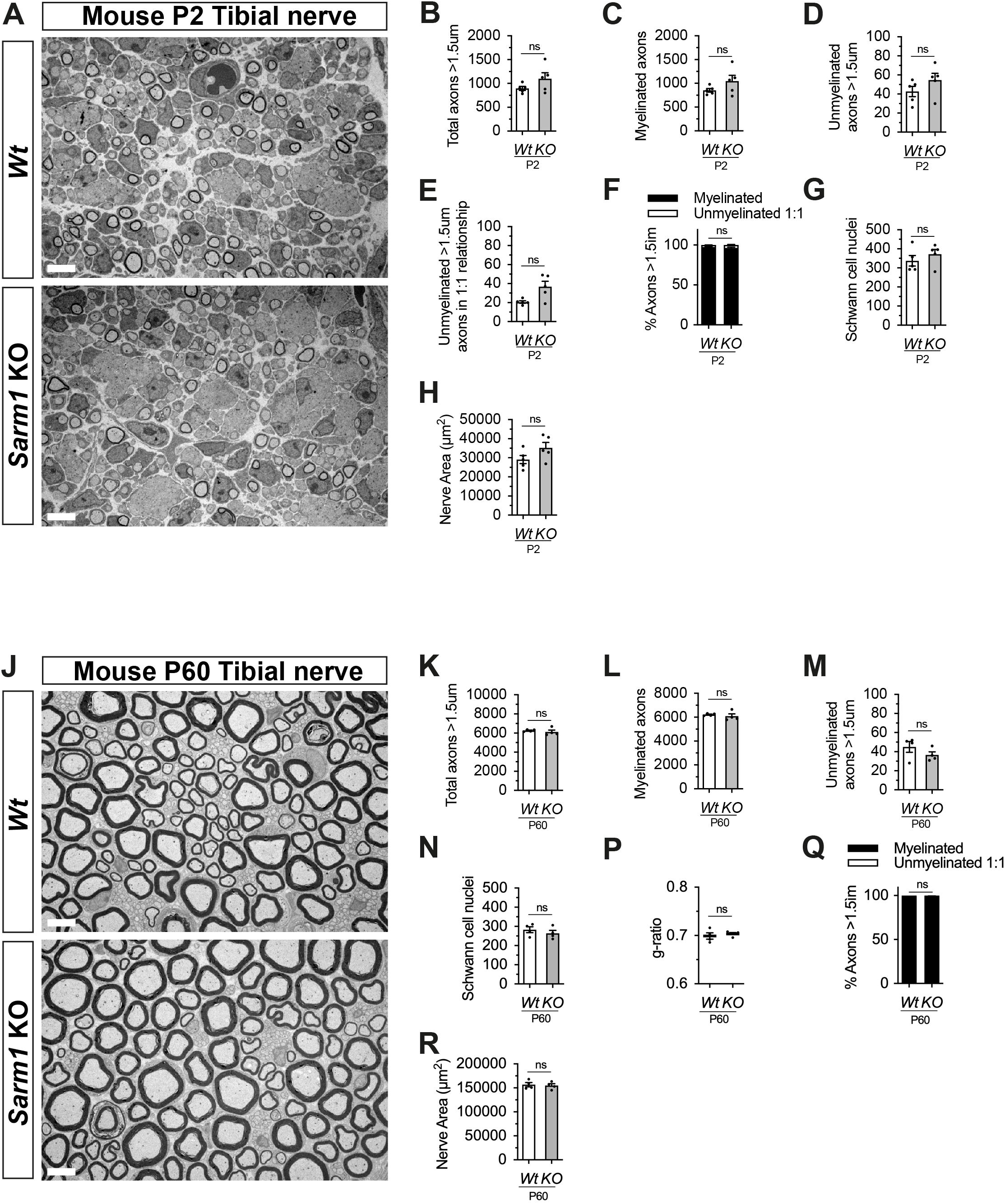
PNS myelination and myelin maintenance are normal in *Sarm1* null mice. A) Representative electron micrographs taken at x3000 magnification from *Wt* and *Sarm1* knockout (KO) tibial nerves at postnatal day 2 (P2). Scale bar 5 μm. B) Per nerve profile, the total number of axons >1.5 μm quantified are not significantly different in *Wt* and *Sarm1* KO nerves (n=5; p=0.3095). C) The number of myelinated axons quantified per nerve profile are similar in both *Wt* and *Sarm1* KO nerves (n=5; p=0.3095). D) The number of unmyelinated axons >1.5 μm is not significantly different in *Wt* and *Sarm1* KO nerves (n=5; p=0.1508). E) Per nerve profile, the number of unmyelinated axons >1.5 μm in a ratio 1:1 is slightly higher in *Sarm1* KO nerves compared to *Wt*; however, this does not reach significance (n=5; p=0.0794). F) The percentage of myelinated axons versus non-myelinated axons present in *Wt* and *Sarm1* KO nerves is not significantly different (n=5; p=0.1508). G) The number of Schwann cell nuclei quantified per nerve profile is not significantly different between *Wt* and *Sarm1* KO nerves (n=5; p=0.5). H) The mean total nerve area of *Wt* (29131 μm^2^) and *Sarm1* KO (35291 μm^2^) nerves is not significantly different (n=5; p=0.1508). J) Representative electron micrographs taken at x3000 magnification from adult *Wt* and *Sarm1* KO tibial nerves at P60. There are no ultrastructural differences in the tibial nerves of adult *Sarm1* KO compared to those of WT tibial nerves. Scale bar 5 μm. K) Per nerve profile, the total number of axons >1.5 μm quantified is similar in *Wt* and *Sarm1* KO nerves (n=4; p=0.3429). L) The number of myelinated axons present in *Wt* and *Sarm1* KO nerves is not significantly different (n=4; p=0.3429). M) Per nerve profile, the number of unmyelinated axons >1.5 μm is not significantly different in *Wt* and *Sarm1* KO nerves (n=4; p=0.3429). N) The number of Schwann cell nuclei is similar in both *Wt* and *Sarm1* KO nerves (n=4; p=0.4857). P). Myelin sheath thickness as depicted by g-ratios is similar in both *Wt* and *Sarm1* KO nerves (n=4; p=0.4857). Q) The percentage of myelinated versus non-myelinated axons present in both *Wt* and *Sarm1* KO nerves is not different (n=4; p=0.3429). R) Per nerve profile, the total nerve area of both *Wt* (156715 μm^2^) and *Sarm1* KO (154927 μm^2^) nerves is very similar (n=4; p=0.8857).

We then looked at adult (P60) tibial nerves by EM to investigate whether myelin maintenance is affected by loss of *Sarm1* in the PNS. Interestingly we no longer saw a trend towards increased axon numbers in the *Sarm1* KO at P60 (Fig.5J-M). Furthermore, the proportion of unmyelinated and myelinated axons, Schwann cell nuclei as well as myelin sheath thickness, measured by g-ratio and nerve area were all similar between *Wt* and *Sarm1* KO samples (Fig.5N-R).

To test whether there were any gene expression differences between *Wt* and *Sarm1* KO tibial nerves we looked at gene and protein expression by quantitative reverse transcriptase polymerase chain reaction (qPCR) and western blot respectively. *Sarm1* KO nerves had no differences in myelin gene or other Schwann cell-specific gene expression (*Cdh1, Egr2, Mbp, Mpz, Sox10*), immature or Schwann cell injury gene expression (*Fos, Jun, Sox2*) or cytokine or chemokine expression (*Ccl2, Ccl3, Ccl4, Ccl5, Il1b, IL6, IL10*). We did however detect that *Sarm1* KO tibial nerves expressed a two-fold higher level of the pro-apoptotic gene, *Xaf1*, which has also been reported to be upregulated in mouse brain and macrophages from this particular *Sarm1* KO mouse line (Fig.6A; Uccellini et al., 2020; Zhu et al., 2019). Additionally, we found no differences in protein levels for EGR2/KROX-20, JUN, and myelin proteins, myelin protein zero (MPZ) and MBP (Fig.6B-G).

**Figure 6.**
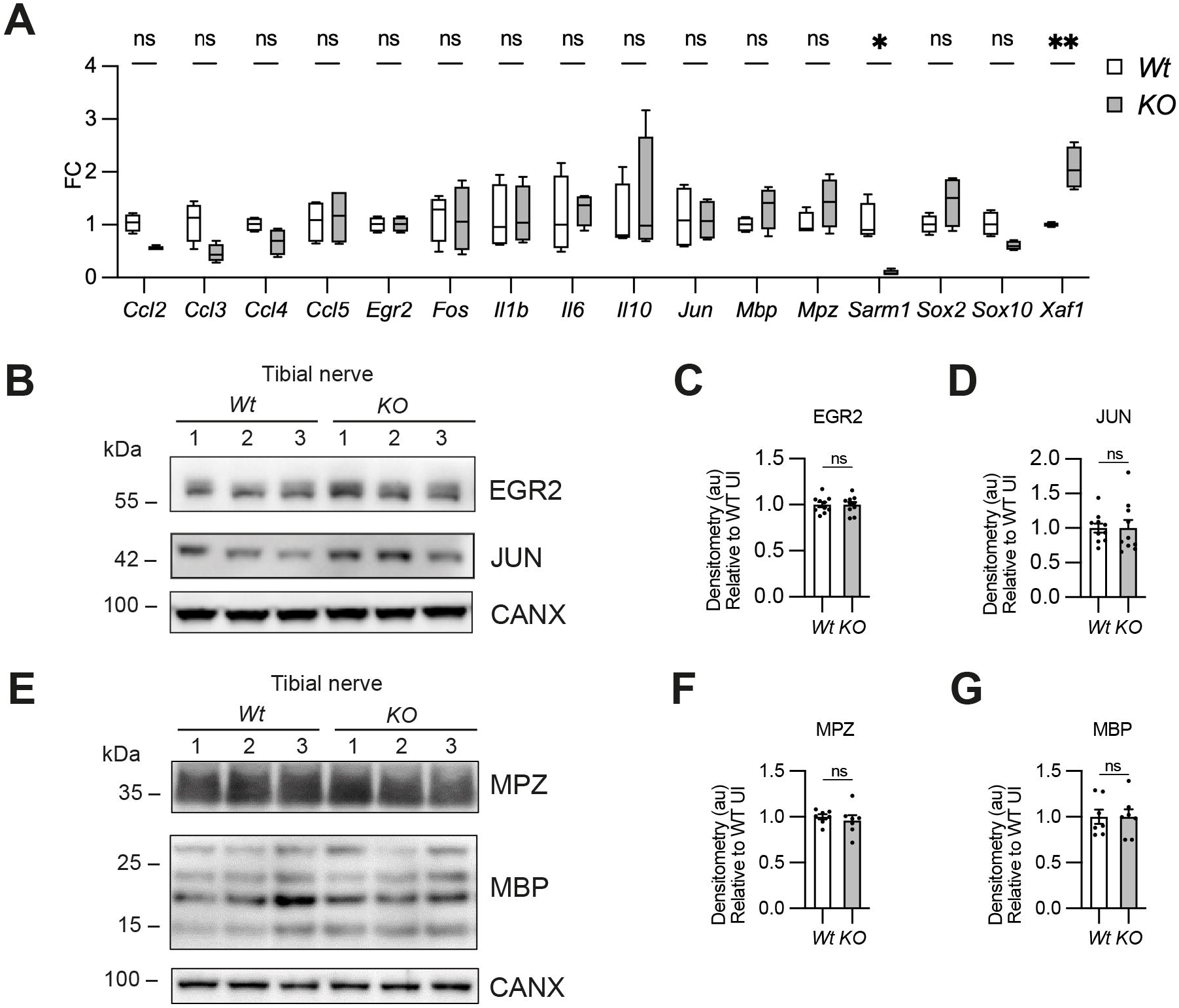
Myelin gene expression is normal in *Sarm1* null peripheral nerves. A) Relative mRNA expression for chemokine, Schwann cell injury and myelin genes in the P60 uninjured tibial nerve of *Wt* and *Sarm1* KO mice. All fold change values normalized to uninjured *Wt* tibial nerve (n=4; *p<0.05, **p<0.01). B) Representative Western blot image of tibial nerve protein extracts from P60 *Wt* and *Sarm1* KO mice. The image shows no difference in levels of EGR2 and JUN, between *Wt* and *Sarm1* KO nerves. C) There is no significant difference in EGR2 levels in *Wt* and *Sarm1* KO nerves (n=7; p=0.9118). EGR2 protein levels are normalized to the levels in *Wt* nerves, which are set as 1. D) There is no significant difference in JUN levels in *Wt* and *Sarm1* KO nerves (n=7; p=0.5787). The quantifications are normalized to the levels in *Wt* nerves, which are set as 1. E) Representative Western blot image showing no difference in levels of Myelin protein zero (MPZ) and Myelin basic protein (MBP), between P60 *Wt* and *Sarm1* KO nerves. F) There is no significant difference in MPZ levels in *Wt* and *Sarm1* KO nerves (WT n=6; KO n=5; p=0.3176). The quantifications are normalized to the levels in *Wt* nerves, which are set as 1. G) There is no significant difference in MBP levels in *Wt* and *Sarm1* KO nerves (n=7; p>0.9999). The quantifications are normalized to the levels in *Wt* nerves, which are set as 1.

Thus, myelination and myelin maintenance are unperturbed and myelin gene and protein expression are normal in the PNS in the absence of *Sarm1* in the mouse.

### CNS myelination and myelin gene expression are normal in the *Sarm1* KO mouse

We have shown that oligodendrocytes in the dorsal and ventral spinal cord of *Sarm1* mutant zebrafish larvae myelinate normally. To confirm whether CNS myelination is also unaffected in *Sarm1* KO mice, we assessed adult P60 optic nerves by EM. We found that *Sarm1* KO optic nerves were morphologically indistinct from *Wt* samples, with similar numbers of total axons, ratio of myelinated to unmyelinated axons and a similar cross-sectional nerve area (Fig.7A-E). Furthermore, by qPCR, *Sarm1* KO optic nerves showed no differences in gene expression for oligodendrocyte/myelin markers (*Mbp, Olig2, Plp1, Sox10*), cytokine or chemokine expression (*Ccl2, Ccl3, Ccl4, Ccl5, IL1b, IL6*), apart from an upregulation of *IL10* (Fig.7F). Interestingly although there was a significant difference in *Sarm1* mRNA expression between *WT* and KO samples, as one would expect, we did not detect any upregulation of *Xaf1* in the optic nerve (Fig.7F). We also tested MBP protein levels by western blot in *Sarm1* KO optic nerves and found no differences compared to *WT* nerves (Fig.7G and H).

**Figure 7.**
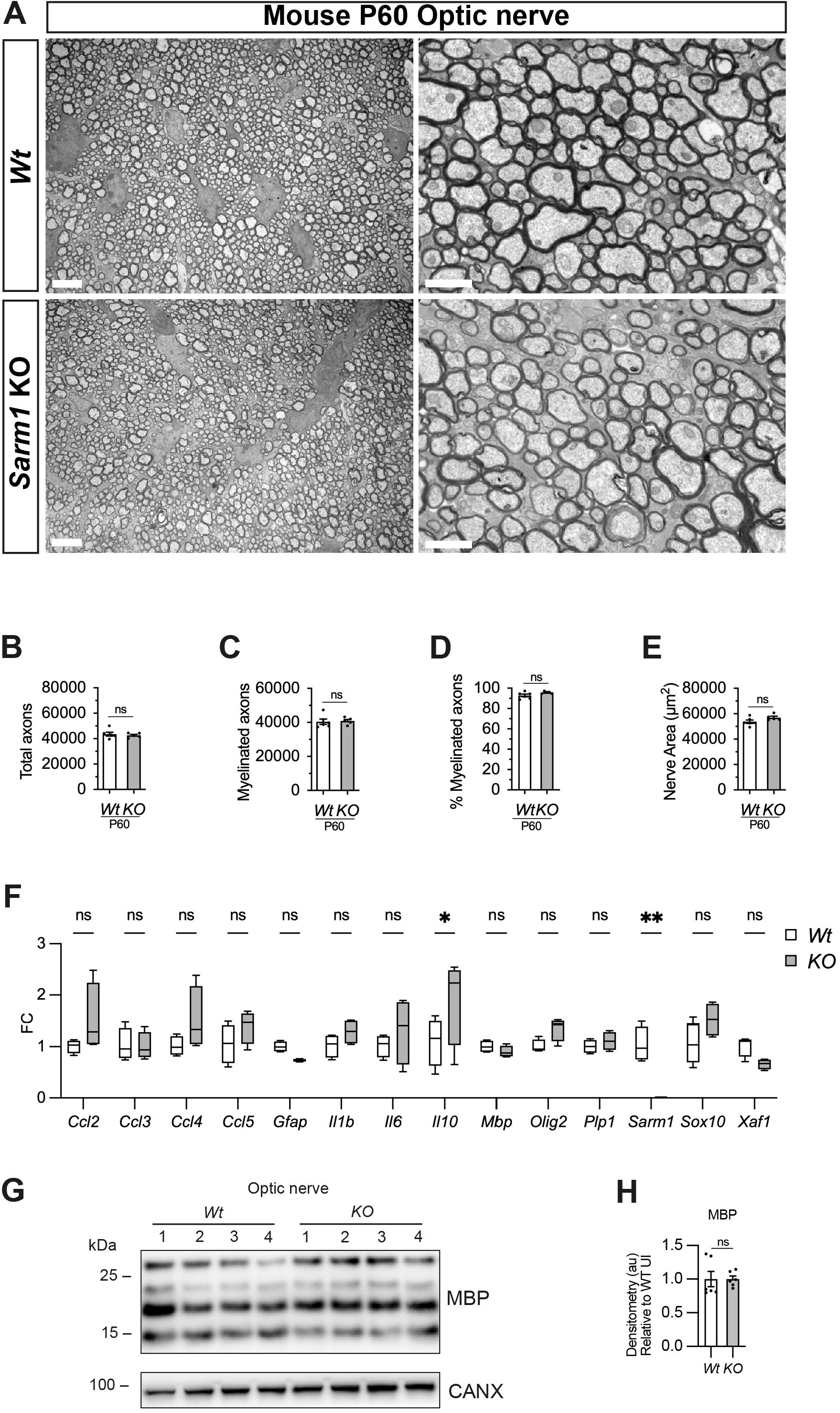
Optic nerve myelination is normal in *Sarm1* null mice. A) Representative electron micrographs taken at x3000 (left hand side panels) and x12000 (right hand side panels) magnification from adult *Wt* and *Sarm1* KO optic nerves at postnatal day 60 (P60). There are no ultrastructural differences in the optic nerves of *Sarm1* KO compared to those of *Wt* animals. Scale bar 5 μm (x3000) and 2 μm (x12000). B) The total number of axons quantified per optic nerve profile are not significantly different between *Wt* and *Sarm1* KO (n=6 WT; n=5 KO; p=0.9307). C) The number of myelinated axons quantified in *Wt* and *Sarm1* KO optic nerves is similar (n=6 WT; n=5 KO; p=0.4286). D) The percentage of myelinated axons quantified in *Wt* and *Sarm1* KO optic nerves is not significantly different (n=6 WT; n=5 KO; p=0.0736). E) Per nerve profile, the total nerve area of both *Wt* (53686 μm^2^) and *Sarm1* KO (56979 μm^2^) optic nerves were not significantly different (n=6 WT; n=5 KO; p=0.1255). F) Relative mRNA expression for chemokine, oligodendrocyte and myelin genes in the P60 uninjured optic nerve of *Wt* and *Sarm1* KO mice. All fold change values normalized to uninjured WT optic nerve (n=4; *p<0.05, **p<0.01). G) Representative western blot image of optic nerve protein extracts shows no difference in levels of MBP, between *Wt* and *Sarm1* KO nerves. H) There is no significant difference in MBP expression between *Wt* and *Sarm1* KO optic nerves (n=6; p=0.3939). The quantifications are normalized to the levels in uninjured *Wt* nerves, which are set as 1.

In conclusion there are no observable defects in myelin in adult *Sarm1* KO mouse optic nerves.

## Discussion

A number of studies investigating the role of SARM1 in the nervous system have documented that PNS and CNS myelin appears normal in the adult *Sarm1 KO* mouse however there have been no in-depth quantitative studies of both PNS and CNS myelination to confirm or refute this (Osterloh et al., 2012; Geisler et al., 2016; Marion et al., 2019; Ko et al., 2020). Additionally, there has been one study that generated a Crispr-Cas9 *sarm1* mutant zebrafish however they did not comment on whether PNS or CNS myelination was normal (Tian et al., 2020). Furthermore, there have been no studies investigating whether SARM1 is present in PNS and CNS myelinating glia.

SARM1, unlike other members of the MyD88 family, is abundant in the murine nervous system, particularly in neurons (Kim et al., 2007; Chen et al., 2011). SARM1 protein is not found in mouse microglia but is expressed in astrocytes and plays a role in neuroinflammation (Lin et al., 2014; Liu et al., 2021; Jin et al., 2022). We have shown that SARM1 is present in mouse oligodendrocytes *in vitro* and *in vivo* and zebrafish oligodendrocytes express *sarm1* mRNA. In contrast, Schwann cells and satellite glia do not contain detectable SARM1 protein and *sarm1*/*Sarm1* mRNA is found at very low levels in Schwann cells in developing zebrafish larvae and in mouse cell culture. Additionally, cultured mouse Schwann cells are insensitive to application of high dose specific SARM1 activators, whereas cultured oligodendrocytes undergo cell death. We have also shown that in the absence of Sarm1/SARM1, PNS and CNS myelination proceeds normally in both zebrafish and mice and that SARM1 does not play a role in myelin maintenance in the murine PNS or CNS. This adds to previous findings that conduction velocities measured by neurophysiological investigation of adult *Sarm1* KO mouse sciatic nerves were similar to control mice and that *Sarm1* KO adult mice have normal myelinated axon numbers in the corpus callosum (Geisler et al., 2016; Marion et al., 2019).

In the PNS, in addition to Schwann cells, we have also shown that satellite glia, do not appear to have identifiable levels of SARM1 protein. SARM1 is also unlikely to be present at high levels in PNS resident macrophages since studies on other peripheral macrophage populations found relatively low levels of SARM1 in these cells and no effect of *Sarm1* deletion on macrophage function or gene expression (Kim et al., 2007; Uccellini et al., 2020). It has been postulated that one of the purposes of SARM1, a toll like adapter protein and remnant of the innate immune system, and the wider axon degeneration machinery is to bring about compartmentalized neurodegeneration to prevent spread of viral pathogens throughout the nervous system (Tsunoda, 2008). In mouse CNS myelinating cocultures, loss of Sarm1 protects neuronal somas against infection and cell death (Crawford et al., 2022). Zika virus preferentially infects oligodendroglia and astroglia in neuron/glia cocultures and causes glia cell death (Cumberworth et al., 2017). Activation of SARM1 is known to cause neuronal cell body death independently of axonal degeneration and in this study we have shown that it can cause oligodendroglial death (Sasaki et al., 2020; Loreto et al., 2021). It is currently unknown if CNS glial cells upregulate *Sarm1* expression in response to zika infection and whether the glial cell death is SARM1 dependent? Additionally, oligodendrocyte death is curbed in the *Sarm1* KO mouse in a neuroinflammatory model of glaucoma. The authors postulate that the death is likely secondary to the role of Sarm1 in axons yet there remains the intriguing possibility that *Sarm1* may, in addition, be required cell-autonomously in oligodendrocytes for some or all of the cell death in this model (Ko et al., 2020). Interestingly, astrocytes do appear to upregulate expression of SARM1 in response to spinal cord injury and also to modulate neuroinflammation in experimental autoimmune encephalomyelitis (Liu et al., 2021; Jin et al., 2022). Finally, while we have investigated two commonly used vertebrate animal models in this study, it will be important in the future to investigate *SARM1* expression and function in human PNS and CNS glia.

In summary, our findings suggest that using *Sarm1* mutant mice and zebrafish to model axonopathy in various disease models is a viable approach as myelination is unlikely to be perturbed by *Sarm1* deletion. Additionally, our data remains consistent with the view that targeted knockdown or inhibition of SARM1 function is a promising strategy to ameliorate various neurological diseases, such as ALS or chemotherapy induced neuropathy, as this is unlikely to aberrantly affect PNS or CNS myelin in humans.

## Acknowledgements

We thank David Lyons for the *Tg[MBP:eGFPCAAX]* zebrafish and Yi-Ping Hseuh for the rabbit polyclonal anti-SARM1 antibody. We thank members of the Coleman, Steventon and Monk labs for useful discussions. We thank Jiakun Chen for assistance with *Sarm1* mutant zebrafish and Kevin O’Halloran and the team at the Cambridge Centre for Advance Imaging for assistance with microscopy. We thank Guillermina López-Bendito, Isabel Pérez-Otaño and Ueli Suter for transgenic mouse lines and David Lyons for transgenic zebrafish lines. Clara Mutschler was funded by a Medical Research Council (UK) studentship (2251399). Peter Arthur-Farraj (206634/Z/17/Z), Andrea Loreto (210904/Z/18/Z) Roger Barker (203151/Z/16/Z) and Michael Coleman (220906/Z/20/Z) were funded by the Wellcome Trust (UK). Ben Steventon was supported by a Henry Dale Fellowship jointly funded by the Wellcome Trust and the Royal Society (109408/Z/15/Z). Kelly Monk was funded by the National Institute of Neurological Disorders and Stroke Awards (R01NS079445). Jose A. Gomez-Sanchez was funded by a Miguel Servet Fellowship (CP22/00078) from the Instituto de Salud Carlos III. Hugo Cabedo was funded by the Spanish “Ministerio de Economica y Competitividad” (BFU2016-75864R and PID2019-109762RB-I00), ISABIAL (UGP18-257 and UGP-2019-128) and Generalitat Valenciana (PROMETEO 2018/114). Yi-Ping Hsueh and Chiung-Ya Chen were funded by Academia Sinica, AS-IA-106-L04 to Yi-Ping Hsueh. For the purpose of Open Access, the author has applied a CC BY public copyright licence to any Author Accepted Manuscript version arising from this submission.

## Materials and Methods

### Animals

All zebrafish research complied with the Animals (Scientific Procedures) Act 1986 and the University of Cambridge Animal Welfare and Ethical Review Body (AWERB), project license code P98A03BF9. Zebrafish (Danio rerio) were maintained at 28°C. *Tg[mbp:eGFPCAAX]* fish were a gift from Dave Lyons (Almeida et al., 2011), and Sarm1^11193^ mutant fish were obtained from ZIRC (Kettleborough et al., 2013). *Tg[mbp:eGFPCAAX]* and *sarm1*^*SA11193*^ embryos were obtained by natural spawning and raised in E3 medium (5 mM NaCl, 0.17 mM KCl, 0.33 mM CaCl2, 0.3 mM MgSO4, and 0.1% methylene blue) in Petri dishes until 24 hours post fertilization. Embryos were then incubated in E3 medium supplemented with 0.003% phenylthiourea (PTU) to prevent pigmentation. F2 generation fish were used for analysis. *sarm1*^*+/+*^ (*wt*) were compared to *sarm1*^*SA11193/*SA11193^ (*sarm1* mutant) for all experiments. Mouse research complied with European Union guidelines and protocols were approved by the Comité de Bioética y Bioseguridad del Instituto de Neurociencias de Alicante, Universidad Miguel Hernández de Elche and Consejo Superior de Investigaciones Científicas (http://in.umh-csic.es/), Reference number 2017/VSC/PEA/00022 tipo 2. *Sarm1* knockout mice on C57BL/6J background (Kim et al., 2007) were bred from heterozygote crosses and F1 littermates were used for analysis. Mice were genotyped as previously described (Kim et al., 2007). *Tg[Plp1-cre/ERT2]* mice (MGI:2663093), expressing Cre recombinase under a tamoxifen inducible Proteolipid 1 (*Plp1*) promoter; *Tg[Sox10-cre]* mice (MGI:3586900), expressing Cre recombinase under control of the *Sox10* promoter; and *B6*.*Cg-Gt(ROSA)26Sor*^*tm14(CAG-tdTomato)Hze*^*/J* (MGI:3813512) that express tdTomato upon removal of a floxed stop codon under control of the *Rosa26* locus were all obtained from Jackson Laboratories (Leone et al., 2003; Matsuoka et al., 2005; Madisen et al., 2010). *Tg[Plp1-cre/ERT2]* mice received an injection of 100 mg/kg tamoxifen during 5 consecutive days at P45. Tamoxifen (Sigma-Aldrich) was dissolved in corn oil (Sigma-Aldrich) and absolute ethanol (Merck) (10:1). Nerves were fresh frozen after two weeks of first tamoxifen injection.

### Genotyping zebrafish

Zebrafish genomic DNA was isolated from adult zebrafish fins and incubated at 55° C overnight with 0.5mg/ml Proteinase K in a 10mM Tris-HCL buffer containing 50mM KCL and 10% Tween20 and 10% NP40. PCR was run using primers, Forward: 5’-TCTGGAGCTGGTGGAGCCCT-3’; Reverse: 5’-AGTCTAGTTTCTGCCTGACCTTGG-3’. PCR product was then restriction enzyme digested at 37° C for 1 hour with MseI (New England Biolabs) and then run on a 1.5% Agarose gel. WT PCR product produces a band of 233 base pairs. Mutant PCR product produces two bands of 165 and 68 base pairs.

### Plasmids and microinjection

To generate the *neuroD:tdTomato* construct, the p-5E neurod5kb promotor entry vector (Mo and Nicolson, 2011), gifted by Dr A.Nechiporuk. was inserted along with pME tdTomato (Oehlers et al., 2015), gifted by Dr D. Tobin (Addgene plasmid #135202) and p3EpolyA into pDestTol2pA2 using the Gateway™ system as previously described (Kwan et al., 2007). To visualize posterior lateral line neurons, 20pg of the *neuroD:tdTomato* construct were microinjected into one cell stage embryos. Larvae were then fluorescently sorted at 4dpf.

### 2-photon axotomy of PLL axons in larval zebrafish

Axotomies were carried out at the Cambridge Advanced Imaging Centre on a TriM Scope II two-photon Scanning Fluorescence Microscope using Imspector Pro software (LaVision Biotec) and a near-infrared laser source (Insight DeepSee, Spectra-Physics). The laser light was focused by a 25x, 1.05 Numerical Aperture water immersion objective lens (XLPLN25XWMP2, Olympus). Axotomies were carried out by focusing the laser at a 6×6 μm section of the posterior lateral line using 100% laser power.

### Schwann cell culture

Freshly plated mouse Schwann cells were obtained from P2-4 sciatic and brachial nerves. Nerves were digested with trypsin and collagenase, centrifuged and plated on laminin/Poly-L-lysine (PLL) coated glass coverslips in defined medium as previously described (Arthur-Farraj et al., 2011).

### Measurement of intracellular NAD^+^

Schwann cells were directly plated on PLL coated 24 well plates and maintained in DMEM/5% horse serum (HS). Schwann and HEK 293T cells were treated with 100μM vacor, 250μM 3-AP, or vehicle controls, and collected in media at 0h and 72h after treatment. Protein was extracted using Pierce™ IP Lysis Buffer supplemented with cOmplete™ Mini EDTA-free Protease Inhibitor Cocktail, and a Bicinchoninic acid (BCA) assay carried out to determine protein concentration. All samples were diluted to 0.5μg/μL. The NAD-glo assay was performed according to manufacturer’s instruction of the NAD^+^/NADH-Glo™ assay by Promega (G9071). In short, 25μL of sample were incubated with 12.5μL 0.4M HCl at 60°C for 15min, before 12.5μL 0.5M Tris base were added. 10μL NAD^+^ standards or sample were then mixed with 10μL NAD-glo master mix (1ml luciferin detection reagent, 5μl reductase, 5μl reductase substrate, 5μl NAD^+^ cycling enzyme, 25μl NAD^+^ cycling substrate) and incubated at RT for 40min. Luminescence was read using a GloMax® Explorer microplate reader and NAD^+^ concentrations determined relative to a NAD^+^ standard curve.

### Oligodendrocyte cell culture

Sprague-Dawley neonatal (≤p7) rat pups were decapitated following lethal overdose with pentobarbital. The brains were dissected and submerged into Hibernate-A low fluorescence media (Transnetyx Tissue, #HALF). The tissue was cut into 1-mm^3^ pieces, washed in HBSS^-/-^ (Gibco, # 11039047), then spun down at 100 g for 1 min at room temperature. To digest the tissue, the tissue was incubated in 34 U/mL papain (Worthington, #LS003127) and 40 μg/mL DNase I (Sigma, #D5025) in HALF, and incubated on an orbital shaker (55 rpm) for 40 min at 37 °C. To obtain a single-cell suspension, the tissue was triturated in HALF supplemented with 2% B27 and 2□mM sodium pyruvate, first using a 5-ml serological pipette and then three fire-polished glass pipettes of descending diameter. The supernatant containing the cells was filtered through 70-μm strainers into a tube containing 90% isotonic Percoll (GE Healthcare, #17-0891-01, in 10□×□PBS pH 7.2 (Gibco, #70013032). The final volume was topped up with DMEM/F12 (Gibco, #31331028) then inverted several times to yield a homogenous suspension with a final Percoll concentration of 22.5%. The single cell suspension was then separated from tissue and myelin debris by gradient density centrifugation at 800 g (without brakes) for 20 mins at room temperature. The myelin debris and supernatant were aspirated, leaving only the cell pellet, which was then resuspended in HBSS^-/-^ to wash out the Percoll. Subsequently, red blood cell lysis buffer (BD Biosciences, #555899) was used to remove red blood cells. OPCs were isolated by positive selection using 2.5 μg A2B5 (Merck Millipore, #MAB312) primary antibody, followed by 20□μl of rat anti-mouse IgM antibody (Miltenyi, #130-047-302) per brain using the MACS protocol according to the manufacturer’s instructions. To collect the A2B5^+^ fraction, the MACS MS column (Miltenyi, #130-042-201) was removed from the magnetic stand (Miltenyi, #130-042-102) and cells were flushed from the column with 1□ml of prewarmed OPC medium (DMEM F/12 containing N-Acetyl cysteine (60 ug/ml, Sigma, #A9165), human recombinant insulin (10 ug/ml), sodium pyruvate (1 mM, Thermo Fisher, #11360-070), apo-transferrin (50 ug/ml, Sigma, #T2036), putrescine (16.1 ug/ml, Sigma, #P7505), sodium selenite (40 ng/ml, Sigma, #S5261), progesterone (60 ng/ml, Sigma, #P0130), bovine serum albumin (330 μg /ml, Sigma, #A4919). Cells were counted, then seeded at a density of 20,000 cells/cm^2^ on poly-D-lysine (5 μg/ml PDL, Sigma #P6407) coated plates.

Freshly isolated OPCs were cultured in OPC medium containing proliferation factors, b-FGF (30 ng/ml, Peprotech, #100-18B) and PDGF (30ng/ml, Peprotech, #100-13a) then changed into OPC media containing T3 (Triiodothyronine, Sigma, #T2877) for 5-7 days to induce OPC to oligodendrocyte differentiation. Whilst in culture, cells were maintained in a humidified incubator at 37 °C, 5 % CO2, 5 % O2, with media changes every 48 h.

### Quantification of vacor treated oligodendrocyte cultures

Rat oligodendrocyte cultures were fixed at various time points following DMSO and Vacor treatment. These cells were immunofluorescently labelled with DAPI and an antibody to SOX10. Three representative images taken at 20x across cultures at different time points were quantified using Fiji for the presence of SOX10 expression. This experiment was repeated three times and average numbers of SOX10 positive oligodendrocytes counted per frame were represented.

### In situ hybridization chain reaction (HCR)

Embryos were fixed using 4% paraformaldehyde in calcium and magnesium free phosphate buffered saline (PBS) and stored at -20C in 100% MeOH. HCR 3.0 was performed as described in (Choi et al., 2018). In short, 2 pmol HCR probes were hybridized at 37°C overnight in hybridization buffer, followed by repeated washing in wash buffer. Probes were detected using 30pmol fluorescent hairpins in amplification buffer overnight at room temperature followed by washing in 5X SSC 0.001% Tween-20. Samples were finally counterstained using 1ug/ml DAPI and mounted in 80% glycerol in a MatTek glass bottom dish. DNA probes for zebrafish *mbp, sarm1, sox10*, fluorescent hairpins and buffers were purchased from Molecular Instruments, Inc.

### Imaging and Image Analysis

Mouse optic nerves were imaged on a Zeiss LSM 900 with airyscan 2, 63x oil objective. All fish were imaged on a Zeiss LSM 700 confocal microscope with a 40x oil objective. For imaging of living larvae, 4 or 5 days post-fertilization animals were anaesthetized in 0.01% tricaine (w/v) and embedded in 1% low melting point agarose (w/v in E3 media supplemented with 0.003% of PTU). For imaging of *Tg[mbp:eGFPCAAX]* zebrafish, images of the spinal cord and PLLn were obtained at the level of the first, third and seventh motor nerves distal to the urogenital opening. For live imaging after PLLn injury, images were taken every 10 minutes for up to 26 hours and embryos maintained at 28°C. Survival of fish was monitored by observing heartbeat and circulation. Confocal images are displayed as grid-stitched maximum intensity protections produced using Fiji (Preibisch et al., 2009; Schindelin et al., 2012). For fluorescence intensity measurements of *Tg(mbp:eGFPCAAX)* fish, integrated densities were obtained from regions of interest. For *mbp* HCR quantification, images of the PLLn and spinal cord were obtained at the level of the first, third and seventh motor nerves distal to the urogenital opening. For statistics coding of color by the intensity of the *mbp* signal, nuclear segmentation was conducted in Imaris (Bitplane) using a surface mask around the DAPI stain with a surface detail of 0.22 μm, and touching surfaces were split using a seed size of 4 μm. For signal intensity analysis, surfaces for the spinal cord and posterior lateral line were drawn in Imaris (Bitplane) and intensity sum of both *mbp* and DAPI signals determined. Intensity was then normalized by the area of the surface. For all experiments between seven to nine fish were imaged (spinal cord and one PLLn) per genotype.

### Antibodies

Immunofluorescence: SOX10 (R&D Systems, 1:100, AF2864, RRID: AB_442208) donkey anti-goat IgG (H+L) Alexa Fluor 488 (Invitrogen, 1:1000, A11057), Anti-Beta III Tubulin (Sigma Aldrich, 1:1000, AB9354, RRID:AB_570918), NF200 (Abcam, 1:1000, ab72997, RRID: AB_1267598), SARM1 rabbit polyclonal antibodies (kind gift from Professor Hsueh, 1:500) and purchased from Abcam (Ab226930, 1:2000, RRID: AB_2893433), tdTomato (SICGEN, 1:500, RRID: AB_2722750) donkey anti-goat IgG (H+L) Alexa Fluor 488 (Invitrogen, 1:1000, A11057), Cy3 donkey anti-rabbit IgG (H+L) (Jackson Immunoresearch, 1:500, 711-165-152), Donkey anti-Mouse IgG (H+L) Highly Cross-Adsorbed Seco Alexa Fluor 488 (Invitrogen, 1:1000, A-21202), Donkey anti-Chicken IgY, Alexa Fluor 647 (Merck, 1:1000,15389818), DAPI (Thermo scientific, 1:2000, 62248).

Western blot: CALNEXIN (Enzo Life Sciences, 1:1000, ADI-SPA-860-D, RRID: AB_312058), JUN (Cell Signalling Technology, 1:1000, 9165, RRID:AB_2130165), EGR2 (EMD Millipore, 1:500, ABE1374, RRID:AB_2715555), MPZ (Aves Labs, 1:2000, PZO, RRID:AB_2313561), MBP (EMD Millipore, 1:1000, AB9348, RRID: AB_2140366), Anti-mouse IgG HRP-linked antibody (Cell Signaling Technology, 1:2000, 7076S), Anti-rabbit HRP-linked antibody (Cell Signaling Technology, 1:2000, 7074S), Goat pAb to Chicken IgY H+L (HRP) (Abcam, 1:2000, ab97135).

### Immunofluorescence

Sciatic and optic nerves were dissected from *Wt* and *Sarm1* KO mice and directly embedded in O.C.T. compound, without fixing, and stored at -80° C until ready for cryosectioning. Cryosections at 5μm thick were collected onto SuperFrost plus slides. These slides were post-fixed in 4% PFA at room temperature for 10mins followed by three washes of 5mins each with 1xPBS. The slides were then immersed in 100% methanol at -20° C for 10mins, followed by three washes of 5mins each with 1xPBS. For anti-Tdtomato sections were immersed in 100% Acetone for 5mins at room temperature before blocking solution was added. The slides were then blocked in 5% horse serum (HS)/0.2%Triton x-100/PBS for 30mins at room temperature. Primary antibodies were diluted in blocking solution and slides were incubated overnight at 4° C. The following day, the slides were washed once for 5mins with 1XPBS, twice for 5mins with 0.1%tween/PBS and finally once again for 5mins with 1xPBS before adding secondary antibodies. The corresponding secondary antibodies were diluted at 1:500 along with DAPI at 1:2000 for 2hrs at room temperature and kept in the dark. Slides were then washed once for 5mins with 1XPBS, twice for 5mins with 0.1%tween/PBS and finally once again for 5mins with 1xPBS before mounting using Fluorsave (Calbiochem, 345789) and representative images were taken using Leica DMI6000B at 40x, 63x and 100x. The same protocol was followed for immunofluorescence staining in rat oligodendrocyte cultures and representative images were taken at 20x using Leica DMI8.

### Western blot

Homogenates were obtained from sciatic or optic nerves. These lysates were prepared as described previously (Gomez-Sanchez et al., 2015). Samples were homogenized in ice-cold RIPA buffer with Halt™ Protease Inhibitor Cocktail (100X) (ThermoFisher Scientific 78429). After homogenization, the samples were incubated on ice for 30□min for further lysis. The resulting lysate was centrifuged at 12000□rpm. at 4°C, and the supernatant was taken for the Bicinchoninic acid (BCA) assay (ThermoFisher, 23227). The supernatant was diluted in loading buffer and boiled at 95°C for 5□min. For western blot analysis of SARM1, 20μg of protein was loaded, for JUN and EGR2, 15μg of protein was loaded and for analysis of myelin proteins MPZ and MBP, 7.5μg of protein was loaded. Next, the samples were run in SDS-PAGE and transferred onto a PVDF membrane. After blocking the membrane in 5% skimmed milk (diluted in TBS/0.1% Tween) for 1□h, the membrane was incubated overnight at 4°C with primary antibodies listed. Experiments were repeated at least three times with fresh samples and representative photographs are shown. Densitometric quantification was done using ImageJ. Measurements were normalized to the loading control Calnexin or Beta Actin.

### RNA extraction and qPCR

RNA was extracted from P60 tibial or optic nerves using Trizol (Invitrogen, 15596026) as previously described (Arthur-Farraj et al., 2017). Nerves from two mice were pooled together for each biological replicate. The integrity and quantity of RNA was determined using Nanodrop (ThermoFisher Scientific) and Agilent 2100 Bioanalyzers (Agilent Technologies). 500ng of RNA was used per biological replicate for cDNA conversion using QuantiTect reverse transcription kit (Qiagen, 205311). qPCR was run on a BioRad CFX96 using iTaq with SYBR Green (BioRad, 1725124). *Ankrd27* and *Canx* were used as housekeeping genes. Two technical replicates and five biological replicates were run per experiment. Fold change was calculated using the delta CT method. All primers were designed using Primer blast (NCBI)(Ye et al., 2012).

### Primer sequences

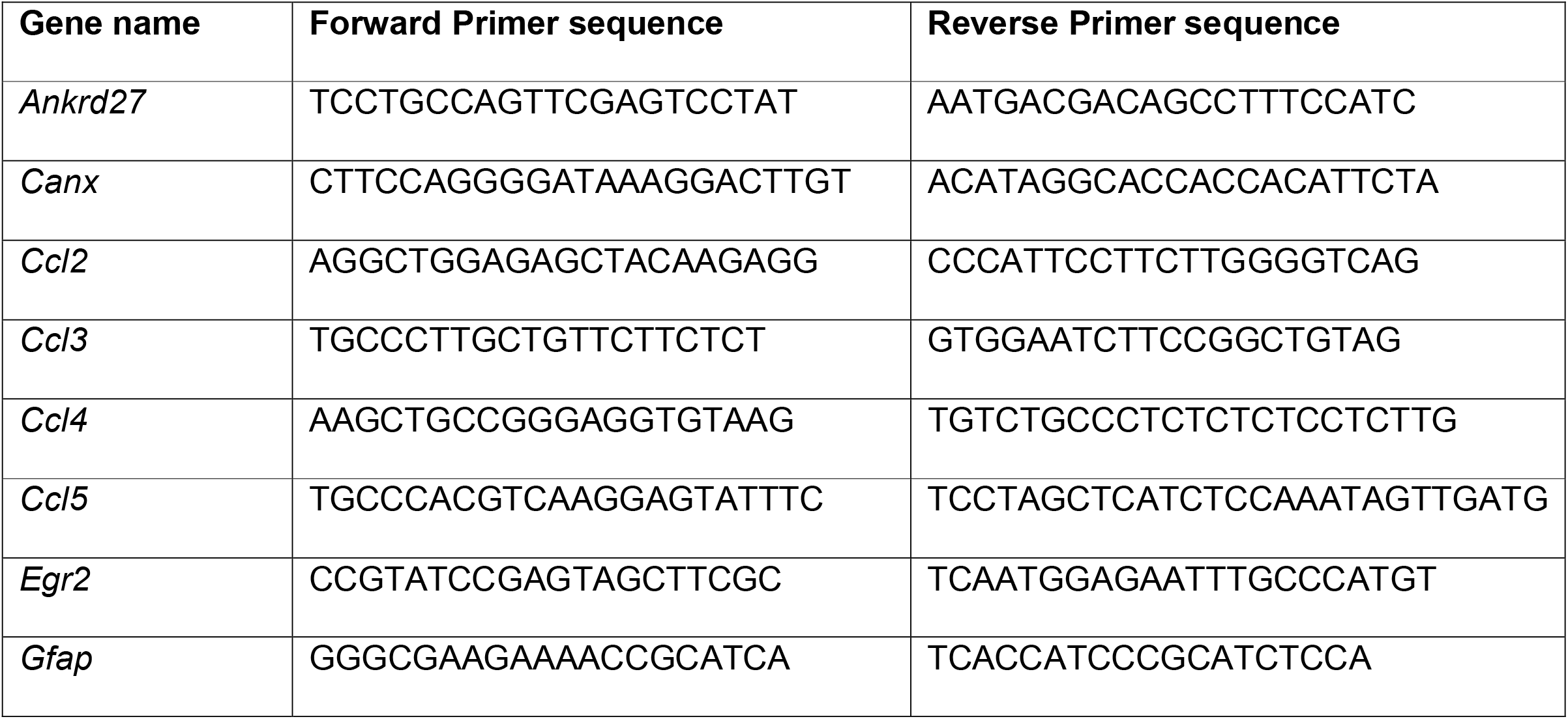
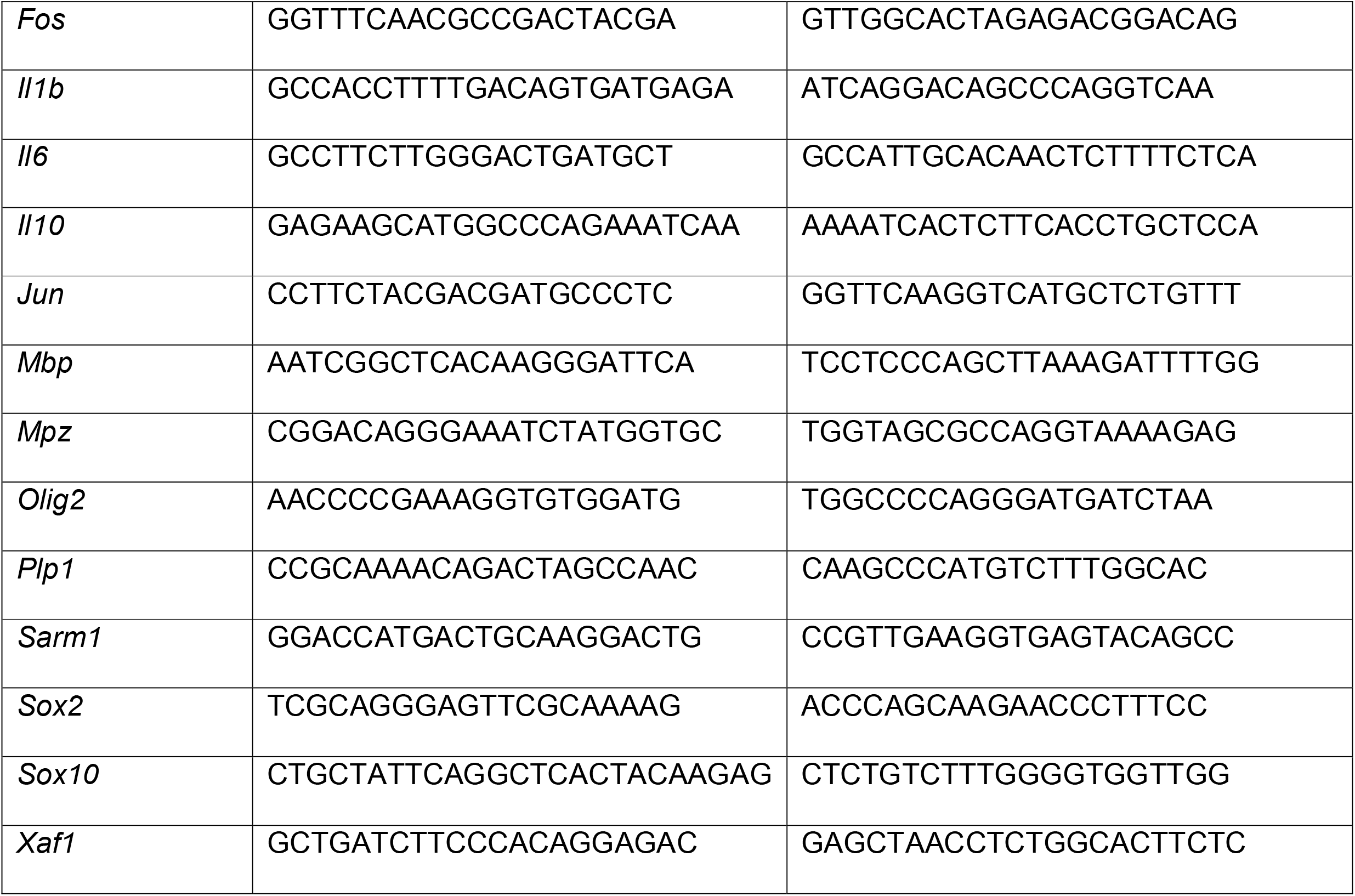

### Electron microscopy

Sciatic and optic nerves were processed as described previously (Gomez-Sanchez et al., 2015). Briefly, samples were fixed in 2.5% glutaraldehyde/2% paraformaldehyde in 0.1 M cacodylate buffer, pH 7.4, overnight at 4°C. Samples were post-fixed with 1% OsO4, embedded in Agar 100 epoxy resin. Transverse ultrathin sections from neonatal (P2) and adult (P60) sciatic nerves (5mm from the notch) or from adult (P60) optic nerves (2mm from the chiasm) were taken and mounted on film.

Photographs were taken using a Jeol 1010 electron microscope with a Gatan camera and software. Images were analyzed using ImageJ. Photographs of sciatic nerves were taken at 3000× magnification to measure the number of myelinated axons, non-myelinated axons bigger than 1.5 μm, and Schwann cell nuclei. The nerve area was measured from photographs taken at 200× magnification. Photographs of optic nerves were taken at 12000x magnification to measure the number of myelinated and non-myelinated axons. The nerve area was measured from photographs taken at 200x magnification. N=5 for all experiments (5 biological replicates: 1 nerve per animal, 5 animals, 15-20 sections per nerve).

### Statistical analysis

Statistical analyses were performed using Graph-Pad Prism software (version 9.1.2). Results are expressed as mean ± SEM. Statistical significance was estimated by Mann-Whitney U-test or unpaired, two tailed students t-test with Bonferonni correction for multiple testing where necessary. P < 0.05 was considered statistically significant. N numbers for all experiments are listed in the figure legends. All quantification was blinded to the assessor.

---

**Supplementary Figure 1. No first layer SARM1 antibody controls**

A) Representative immunofluorescence images of transverse sections of *Wt* and *Sarm1* KO mouse dorsal root ganglia (DRGs) to show lack of SARM1 (blue) signal in the absence of addition of primary antibody. NF-200 (neurofilament light chain; magenta) to show neurons. Scale bar 25 μm. B) Representative immunofluorescence images of transverse sections of *Wt* and *Sarm1* KO mouse dorsal root ganglia (DRGs) to show lack of SARM1 (blue) signal in the KO sample in the presence the primary antibody. The *Wt* panel shows co-localization of NF-200 (magenta) positive neurons with SARM1 (blue). Scale bar 25 μm.

**Supplementary Figure 2. *sarm1* HCR no first layer controls**

A) MAX projection, lateral view of the PLLn of a 5dpf larva with no probes (no first layer control) to *sox10* nor *sarm1* applied. Images show DAPI (gray), and hairpins tagged to Alexa647 (magenta) and Alexa546 (green). Scale bar 25 um. B) MAX projection, lateral view of the spinal cord of a 5dpf larva with no antisense probes to *sox10* nor *sarm1* applied. Images show DAPI (gray), and hairpins tagged to Alexa647 (magenta) and Alexa546 (green). Scale bar 25 μm.

**Supplementary Figure 3. *Sarm1* mRNA but not SARM1 protein is detectable in mouse Schwann cells**.

A) qPCR analysis of *Sarm1* expression in cultured mouse *Wt* and *Sarm1* KO DRGs and *Wt* and *Sarm1* KO Schwann cells (SC, n=4 (4 biological replicates: 4 sciatic nerves from 2 animals pooled per biological replicate); **p<0.01).

B) Freshly plated *Wt* and *Sarm1* KO, P2 dissociated mouse Schwann cells both have undetectable levels of SARM1 protein (blue). Cultures labelled for DAPI (gray) and SOX10 (cyan) (n=3: two-four mice each from three litters producing three separate cultures). Scale bar 50 μm.

**Supplementary Figure 4. Cultured Schwann cells are insensitive to 3-AP induced cell death**. A) Dissociated and freshly plated P2 mouse Schwann cells cultured in water or 250 μM 3-AP for 72 hours show no cell death, judged by propidium iodide (P.I) and DAPI nuclear staining and SOX10 immunocytochemistry (n=3: two-four mice, each from three litters producing three separate cultures). Scale bar 25 μm. B) Intracellular NAD^+^ levels decrease in HEK293 cells when treated for 72 hours with 100 μM vacor. C) Cultured rat oligodendrocytes (SOX10 positive) express SARM1 protein using a commercial polyclonal anti-SARM1 antibody. Zoomed in area (indicated by dashed box). Arrowheads indicate SOX10 positive SARM1 positive cells. Scale bars 50 μm.

**Supplementary Figure 5. Axon degeneration after PLLn laser axotomy is substantially delayed in absence of functional Sarm1 in zebrafish larvae**. A) Schematic of overview of 2-photon laser axotomy of the PLLn (green) in larval zebrafish. B) *wt* and *sarm1*^*SA11193/SA11193*^ (*sarm1 mutant*) fish were injected at the one cell zygote stage with *neuroD:tdtomato* DNA constructs and 2-photon laser axotomy was performed in both genotypes at 4dpf. While *wt* PLLn axons start to degenerate around three hours and are completely degenerated at six. hours post axotomy, PLLn axons in *sarm1 mutant* larvae remain intact at least up until 26 hours post imaging (imaging stopped due to UK home office regulations) (n=5). Scale bar 100 μm.

## Notes

### Competing Interest Statement

The authors have declared no competing interest.

