## Supplementary figures and images for "SARM1 detection in oligodendrocytes but not Schwann cells though *sarm1/Sarm1* deletion does not perturb CNS nor PNS myelination in zebrafish and mice"

### Supplemental Figure 1

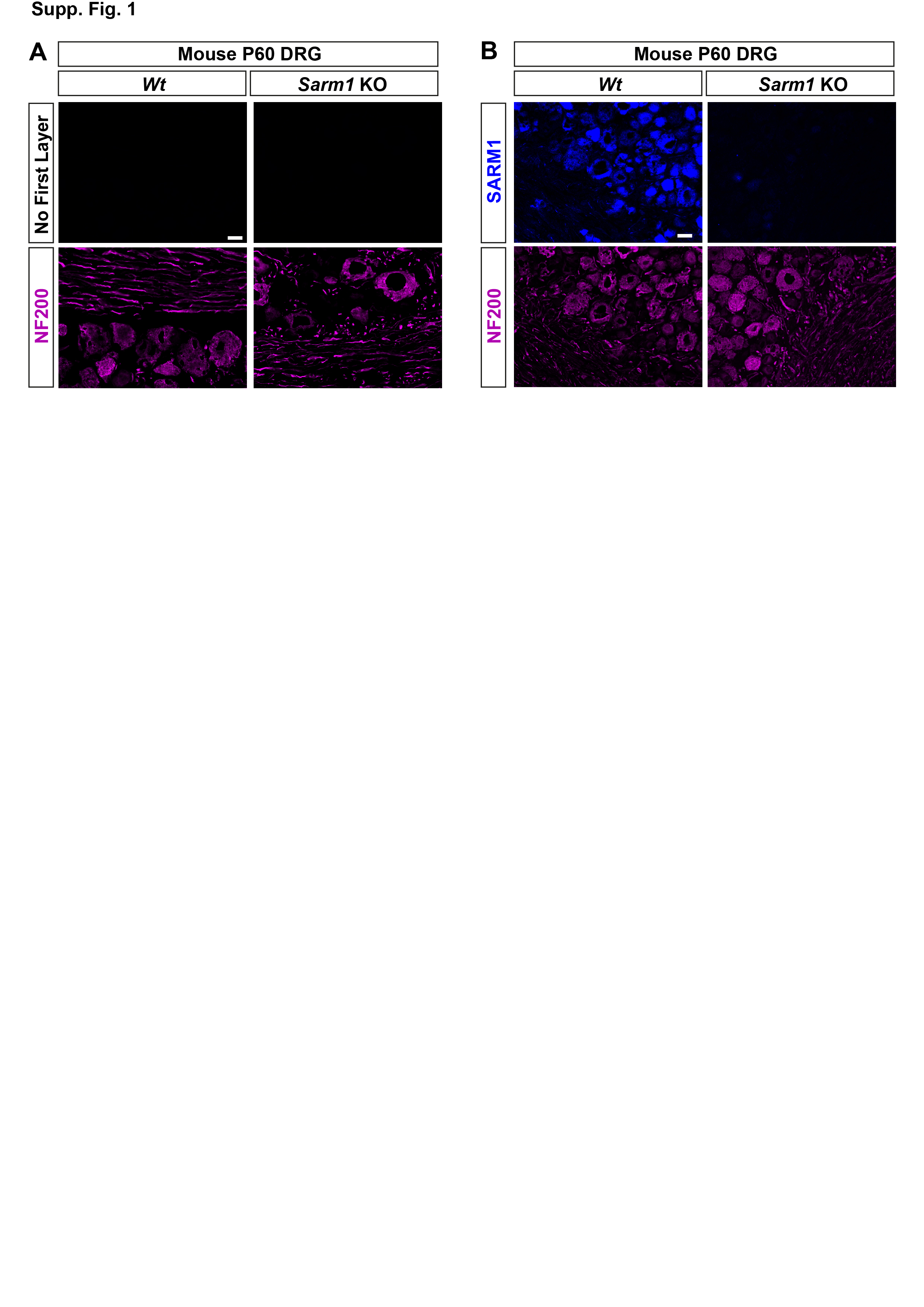

### Supplemental Figure 2

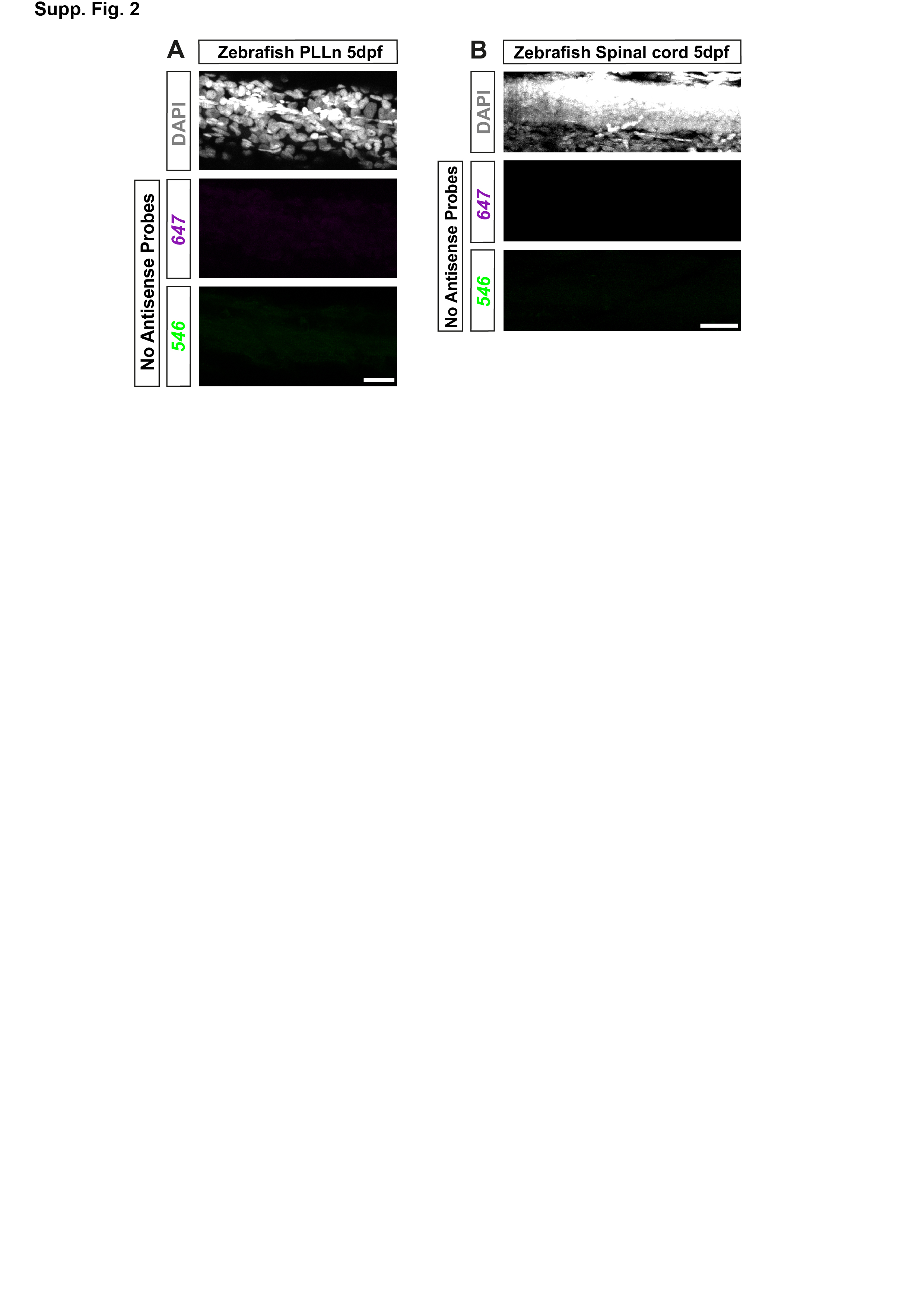

### Supplemental Figure 3

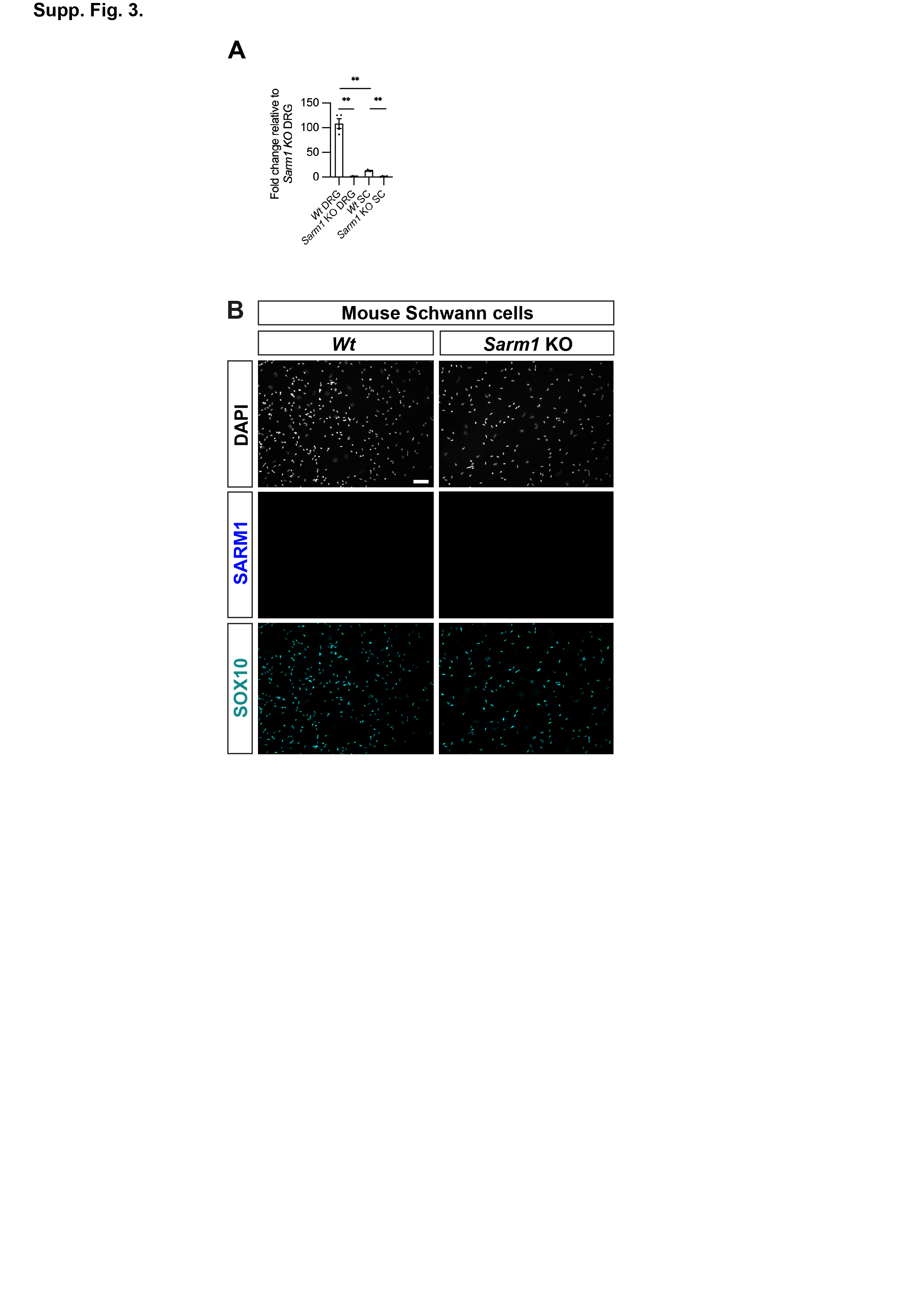

### Supplemental Figure 4

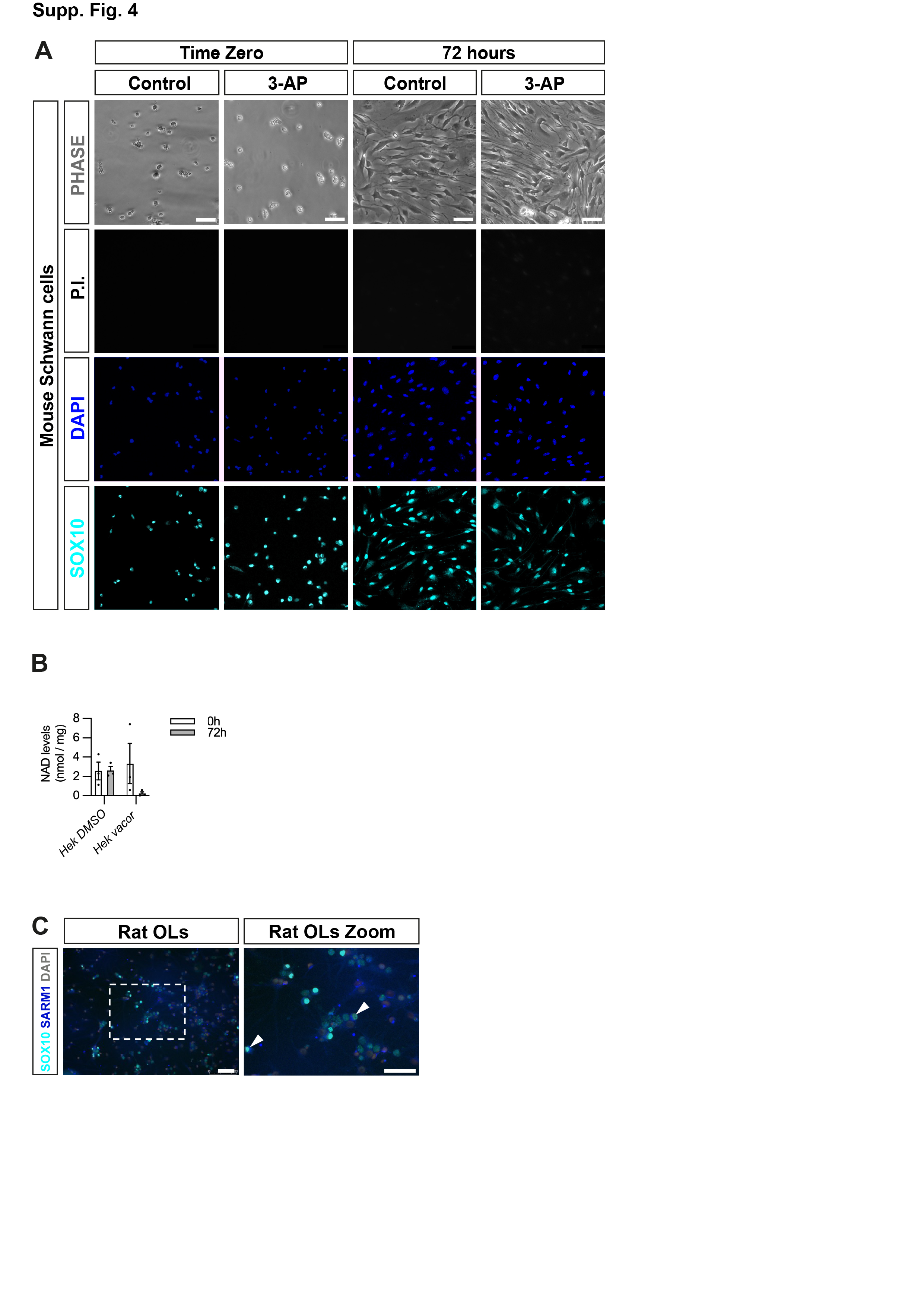

### Supplemental Figure 5

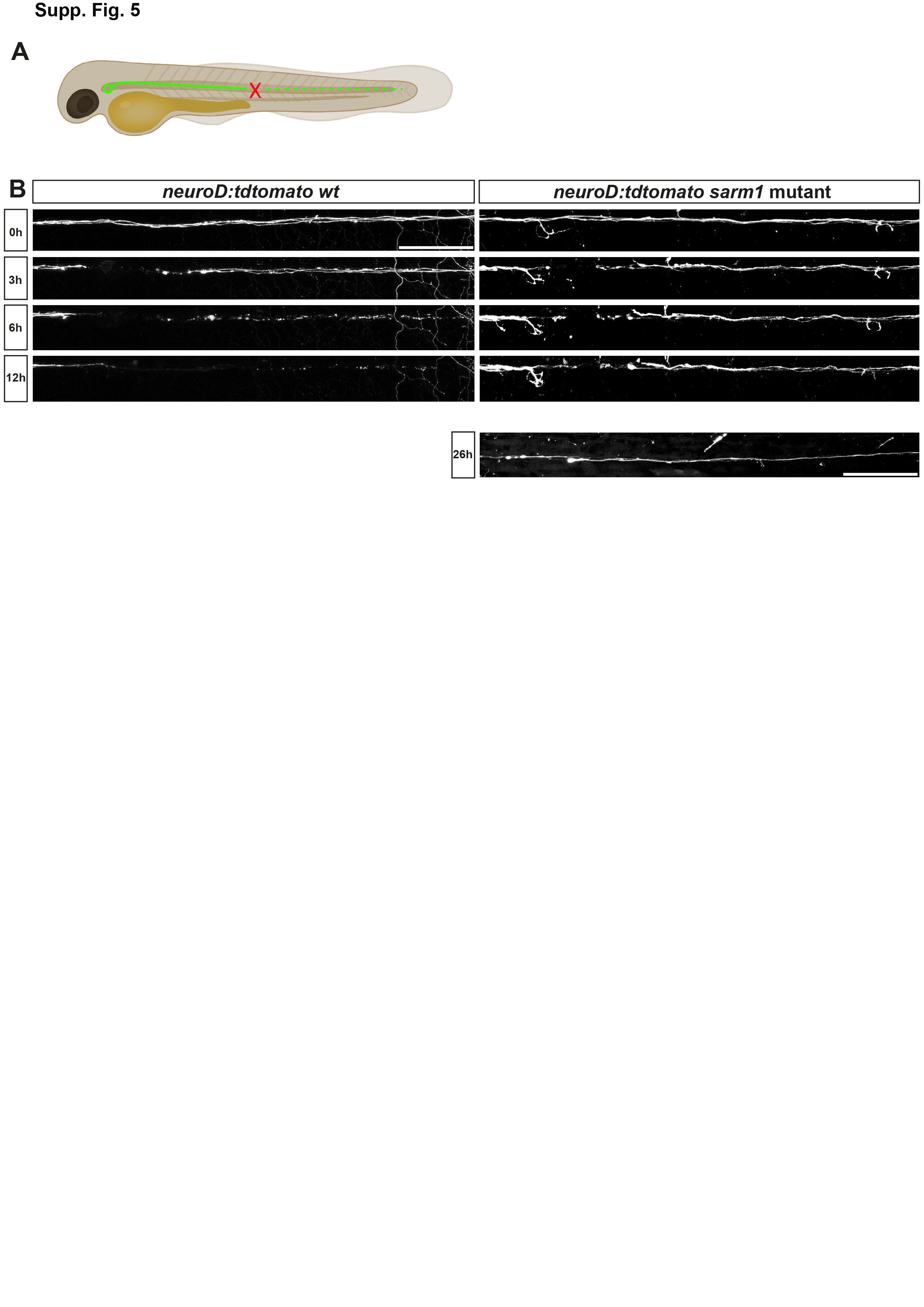
